# Genomic Analysis of Hypoxia Inducible Factor Alpha Evolution in Ray-finned Fishes (Actinopterygii)

**DOI:** 10.1101/2021.04.16.440203

**Authors:** Ian K. Townley, Bernard B. Rees

**Author notes:** Corresponding author: Bernard B. Rees. Significance statement: Vertebrate animals have multiple copies of the hypoxia inducible transcription factor, a critical regulator of oxygen-dependent gene expression, but the number of copies in fishes and their relationships are incompletely understood. This study mines the genomes of ray-finned fishes, finds evidence of gene duplicates not previously appreciated, and clarifies the relationships among duplicates that arose early in vertebrate evolution and those arising from later rounds of genome duplication in fishes.

## Abstract

Two rounds of genome duplication (GD) in the ancestor of vertebrates, followed by additional GD during the evolution of ray-finned fishes (Actinopterygii), expanded certain gene families, including those encoding the hypoxia inducible transcription factor (HIF). The present study analyzed Actinopterygian genomes for duplicates of HIFα, the subunit that confers oxygen-dependent gene regulation. In contrast to tetrapod vertebrates that retain three HIFα genes from the ancestral vertebrate GD, four HIFα forms were found in the genomes of primitive Actinopterygians (spotted gar and Asian arowana). All four forms have been retained in zebrafish and related species (Otocephala) and salmonids and their sister taxa (northern pike) but one of them (HIF4α) was lost during the evolution of more derived fishes (Neoteleostei). In addition, the current analyses confirm that Otocephala retain duplicates of HIF1α and HIF2α from the teleost-specific GD, provide new evidence of salmonid-specific duplicates of HIF1α, HIF2α, and HIF3α, and reveal a broad distribution of a truncated form of HIF2α in salmonids and Neoteleostei. This study delivers a comprehensive view of HIFα evolution in the ray-finned fishes, highlights the need for a consistent nomenclature, and suggests avenues for future research on this critical transcription factor.

## Introduction

Species richness and ecological diversity of the teleost fishes have been attributed, in part, to the evolutionary potential that ensued from multiple genome duplication (GD) events that occurred during the evolution of this group (Volff 2005; Glasauer and Neuhauss 2014; Guo 2017; Ravi and Venkatesh 2018). Two GD occurred at the base of vertebrate evolution (Ohno 1970; Dehal and Boore 2005; Sacerdot et al. 2018), and the evolution of ray-finned fishes (Actinopterygii), which comprise about half of the extant species of vertebrate animals, has been characterized by additional GD: the teleost-specific genome duplication (TGD) occurred early in the evolution of teleost fishes (Postlethwait et al. 2000; Volff 2005; Pasquier et al. 2016; Ravi and Venkatesh 2018) and a more recent GD occurred during the evolution of salmonids (SGD) (Berthelot et al. 2014; Macqueen and Johnston 2014). Although the most common fate of gene duplication is silencing of one copy (nonfunctionalization) (Lynch and Conery 2000; Inoue et al. 2015), a considerable number of duplicates can be maintained by acquiring a new function (neofunctionalization) or by sharing the roles, developmental timing, or tissue specificity of the ancestral gene (subfunctionalization) (Force et al. 1999). Indeed, genomic analyses of teleost fishes shows a high proportion of gene duplicates have been maintained in certain species (e.g., up to 48% in rainbow trout; Berthelot et al. 2014).

The hypoxia-inducible factor (HIF) orchestrates cellular responses to low oxygen in animals (Kaelin and Ratcliffe 2008; Semenza 2011). The active transcription factor is composed of an oxygen-dependent alpha subunit (HIFα) and a constitutively expressed beta subunit (HIFβ), also known as the aryl hydrocarbon receptor nuclear translocator (ARNT), both of which belong to the large family of PAS-domain proteins, so named for conserved domains present in the transcription factors Period, ARNT, and Single-minded (Gu et al. 2000; McIntosh et al. 2010). Tetrapod vertebrates, including humans, have three forms of HIFα encoded by the *HIF1A*, *HIF2A*, and *HIF3A* loci (see Box 1, Supplementary Material online)(Kaelin and Ratcliffe 2008; Semenza 2011; Duan 2016). Previous analyses of HIFα evolution among fishes suggested that most teleost fishes have three forms, which appear to be orthologs of the forms found in tetrapods (Nikinmaa and Rees 2005; Rytkönen et al. 2011; Rytkönen et al. 2013; Townley et al. 2017). However, zebrafish and relatives (*Cyprinidae*) possess six forms and some non-cyprinid species retain a truncated form of HIF2α (Rytkönen et al. 2013; Graham and Presnell 2017; Townley et al. 2017). Rytkonen et al. (2013, 2014) provided evidence that the six forms in *Cyprinidae* arose from the TGD and were maintained in this lineage by subfunctionalization, whereas they were lost, or dramatically truncated, in other lineages.

Here, we reexamined the evolution of HIFα in ray-finned fishes using recently sequenced genomes, including that of spotted gar (*Lepiosteus oculatus*) to represent a lineage that diverged prior to the TGD (Braasch et al. 2016). Specifically, we sought to determine (1) whether any ray-finned fishes retained the four HIFα genes that arose during the two rounds of GD at the base of vertebrate evolution, (2) whether gene duplicates arising from the TGD are seen in fishes other than the *Cyprinidae*, (3) whether HIFα duplicates from the SGD are present in salmonid genomes, and (4) the phylogenetic relationship and distribution of shortened forms of HIF2α.

## Methods

Actinopterygian genomes at NCBI (https://www.ncbi.nlm.nih.gov) or Ensemble (http://www.ensembl.org/) were searched for HIFα genes through June 2020. The corresponding coding sequences (CDS) were checked to ensure each was complete and, when multiple CDSs were available for a single locus, the longest sequence was retained. The final sequence list was curated to avoid overrepresentation of Neoteleostei and to remove sequences with poor chromosomal or linkage group assignment (supplementary table S1, Supplementary Material online).

Multiple sequence alignments (MSA) were made using MAFFT version 7.123b (Katoh and Standley 2013; Katoh and Standley 2016) implemented through the GUIDENCE2 server with Max-iterate of 20 (http://guidance.tau.ac.il/ver2/). For phylogenetic analysis, we used the MSA with the default column cutoff of below 0.93 (Penn et al. 2010; Sela et al. 2015). Bayesian analyses were performed in BEAST 2 v2.6.1 (https://www.beast2.org/) applying an uncorrelated log normal relaxed molecular clock model (Drummond et al. 2006), a Yule model prior, with 10,000,000 chain-length and 100,000 burn-in (Drummond et al. 2012; Bouckaert et al. 2014). Nucleotide analysis employed a general time reversible codon substitution model allowing for invariants and six gamma categories (GTR+I+G). Amino acid analysis employed a JTT matrix-based model (Jones et al. 1992) allowing for invariants and six gamma categories. The maximum clade credibility tree was selected using TreeAnnotator v2.6.0. Maximum likelihood analyses were performed using MEGAX v10.1.8 (Kumar et al. 2018) with 100 bootstrap replicates (Felsenstein 1985) under the same conditions used in Bayesian analyses. The trees with the highest log likelihood were visualized and edited in FigTree v1.4.4 (http://tree.bio.ed.ac.uk/software/figtree/).

Shared synteny among representative species was assessed by determining the flanking deduced open reading frames (ORF) using the NCBI Graphical Sequence Viewer (v3.38.0). If a putative ORF lacked a clear identification, BLASTP was used to compare the deduced protein sequence against Actinopterygii. A small number of putative ORFs could not be identified and were kept in the analysis as “unknowns”. For genes lacking an abbreviation at NCBI, the gene name was used in a search of UniprotKB (https://www.uniprot.org), and the corresponding abbreviation was used.

The N-and C- terminal oxygen-dependent degradation domains (NODD and CODD) and the C-terminal asparaginyl hydroxylation motif (CEVN) were identified from multiple sequence alignments in Jalview v2.10.5 (Waterhouse et al. 2009). The NODD and CODD included the canonical LxxLAP sequence targeted by prolyl hydroxylases and adjacent residues known to play a role in oxygen-dependent regulation of HIFα (Tarade et al. 2019). Consensus sequences were made using the Skylign server (http://skylign.org/) (Wheeler et al. 2014).

## Results and Discussion

Phylogenetic analyses of 114 putative HIFα homologs from 22 species in 14 orders of Actinopterygii (supplementary table S1, Supplementary Material online) resolved four distinct homology groups (fig. 1). This pattern was strongly supported by Bayesian analyses of nucleotide and deduced amino acid sequences (posterior probabilities ≥ 0.89; supplementary figs. S1, S3, Supplementary Material online), as well as by maximum likelihood (bootstrap values ≥ 0.71 for nucleotide analyses and ≥ 0.79 for amino acid analyses; supplementary figs. S2, S4, Supplementary Material online). The branching patterns of species within each homology group generally reflected the known phylogeny of fishes (Hughes et al. 2018), although the relationship of paralogs varied among analyses (see below).

**Fig. 1.**
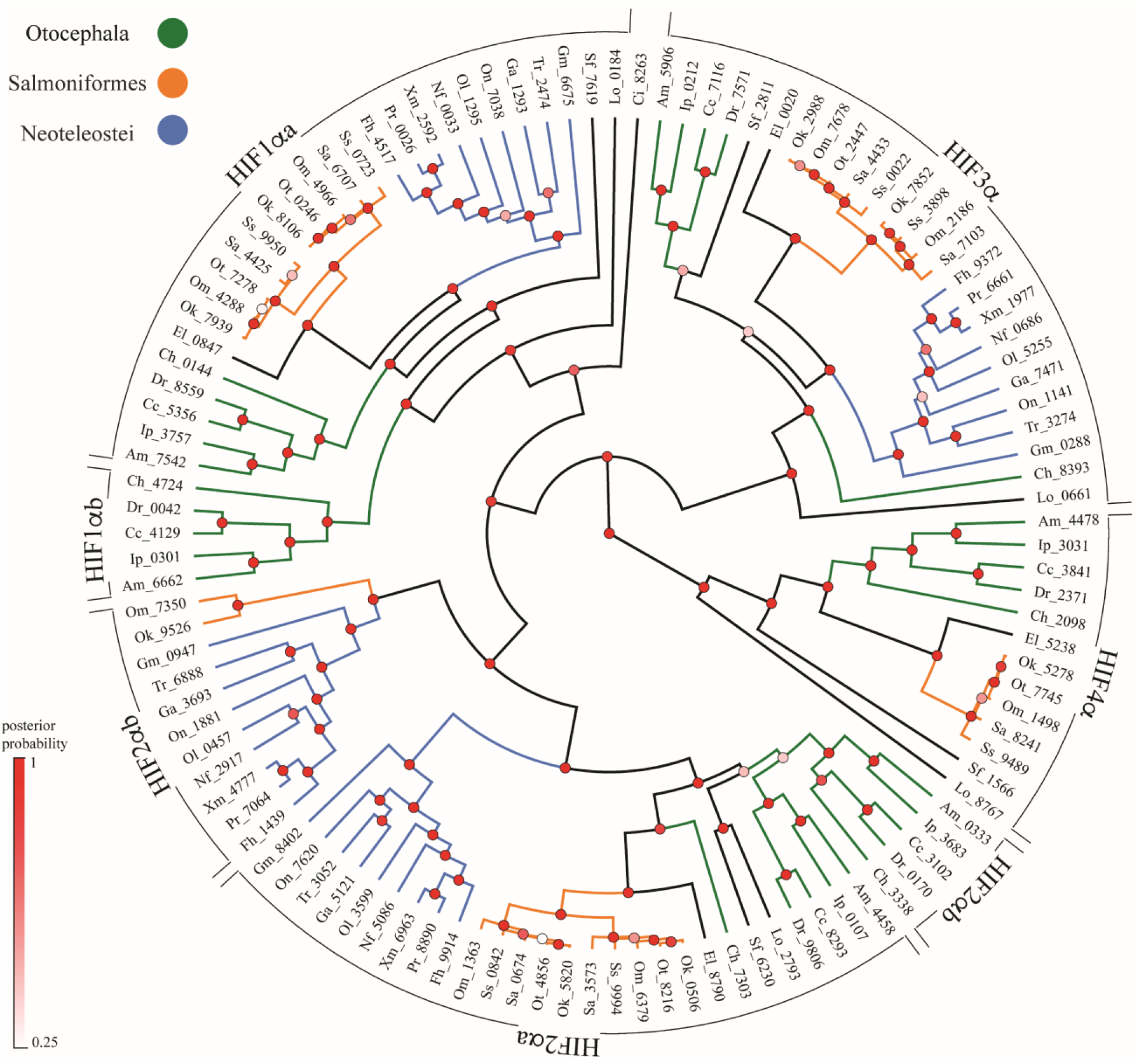
Phylogeny of Actinopterygian HIFα reconstructed by Bayesian inference using full-length coding sequences. Analyses used the General Time Reversible model (GTR) with 6 gamma categories (+G) and allowing for invariants (+I). The tree with maximum clade credibility and mean heights is shown and the nodes are colored by posterior probability values. HIFα homology groups are shown on the periphery and the following taxa are color coded within each group: Otocephala (green); Salmoniformes (orange); Neoteleostei (blue). The outgroup, *Ciona intestinalis*, basal Actinopterygian (spotted gar, *Lepisosteus oculatus*), basal teleost (Asian arowana, *Scleropages formosus*), and sister taxa to Salmoniformes (Northern pike, *Esox lucius*), are not color coded. Sequences are identified by the first letter of the genus and species followed by the last four digits of the NCBI or Ensemble reference gene accession number (see supplementary table S1, Supplementary Material online for a full list of genes).

The spotted gar (*Lepiosteus oculatus)*, a basal Actinopterygian that diverged prior to the TGD, has one homolog in each group. Thus, we infer that the four groups represent products of the two rounds of GD in the ancestor of vertebrates (i.e., ohnologs, Ohno 1970; Wolfe 2000) and hereafter refer to these as HIF1α, HIF2α, HIF3α, and HIF4α. The current results support previous observations (Graham and Presnell 2017) and suggest that several sequences described as “HIFα-like” are HIF3α or HIF4α (fig. 1, supplementary table S1, Supplementary Material online). Like the spotted gar, the Asian arowana (*Scleropages formosus*) has four HIFα genes, one in each of the major groups. The arowana is in the Osteoglossiformes (bonytongues), an ancient order that diverged early from the main branch of teleost evolution (Hughes et al. 2018), but after the TGD (Bian et al. 2016). In this lineage, therefore, one of each pair of HIFα paralogs resulting from the TGD appears to have been lost due to nonfunctionalization, as predicted for other TGD paralogs (Bian et al. 2016).

Our analysis confirms that Cypriniformes (e.g., zebrafish and common carp) have duplicated forms of HIF1α (HIF1αa and HIF1αb) and HIF2α (HIF2αa and HIF2αb), which likely arose during the TGD (Rytkönen et al. 2013). These duplicates are more broadly distributed than previously appreciated, being present in all the species in Otocephala that were examined, including Atlantic herring, Mexican tetra, and channel catfish. More derived fishes appear to lack HIF1αb, although a truncated version of HIF2α has been retained (see below; Rytkönen et al. 2013). Otocephala has two other HIFα forms that group with HIF3α and HIF4α from spotted gar. This placement strongly suggests that these forms represent ohnologs arising from the GD in ancestral vertebrates, rather than HIF3α duplicates arising from the TGD, as previously proposed (Rytkönen et al. 2013).

The current analysis revealed that Salmoniformes have two forms of HIF1α, HIF2α, and HIF3α (fig. 1). These duplicates are not observed in the sister group Esociformes (pike) and the branch lengths joining them are very short, consistent with an origin during the SGD. Berthelot et al. (2014) noted that rainbow trout retained as many as 48% of the gene duplicates, including an over-representation of transcription factors. Because salmonids generally occur in well-oxygenated habitats and have poor hypoxia tolerance, there is no apparent link between HIFα duplication and hypoxia tolerance in this group. Rather, these duplicates may play other roles in salmonid physiology or life-history. Alternatively, duplicates may be destined for nonfunctionalization, a process that is likely still underway in this lineage (Berthelot et al. 2014). Only one salmonid-specific duplicate of HIF4α was recovered in the current analysis, suggesting one duplicate has already been nonfunctionalized. In addition, HIF4α appears to be completely absent in more derived fishes (Neoteleostei).

Genomic analysis also supports the broad distribution of a truncated form of HIF2α, occurring in Salmoniformes and all Neoteleostei examined. The deduced protein sequence corresponds to the N-terminal half of HIF2α, containing the basic helix-loop-helix and PAS domains, which are important for DNA-binding and protein dimerization, but lacking sites for oxygen-dependent regulation of protein stability and gene activation (Rytkönen et al. 2013; Townley et al. 2017). Although speculative, this truncated form of HIF2α might act as a negative regulator by dimerizing with other HIFα subunits and preventing their transcriptional activity, as demonstrated for similarly truncated forms of HIF3α that arise from alternative splicing in mammals (Makino et al. 2002). Alternatively, this form may be undergoing nonfunctionalization, particularly in Salmonifomes in which even shorter forms are present in species not included in this genomic analysis.

Because the relationship of paralogs arising from the TGD based upon sequence analyses alone can be ambiguous (supplementary figs. S1-S4, Supplementary Material online; Parey et al. 2020), we used shared synteny among species representing major fish lineages to clarify the relationships among HIFα paralogs (fig. 2; supplementary table S2, Supplementary Material online). For HIF1α, several flanking genes are conserved from spotted gar (*L. oculatus*) through Neoteleost (represented by *X. maculata*). A subset of up to eight neighboring genes is shared among HIF1αa in Otocephala (represented by *D. rerio*) and HIF1α in other fishes, but not found for Otocephala HIF1αb. This pattern supports the view that HIF1αa of Otocephala is orthologous with HIF1α in other fishes, whereas HIF1αb was lost in those lineages (Rytkonen et al. 2013).

**Fig. 2.**
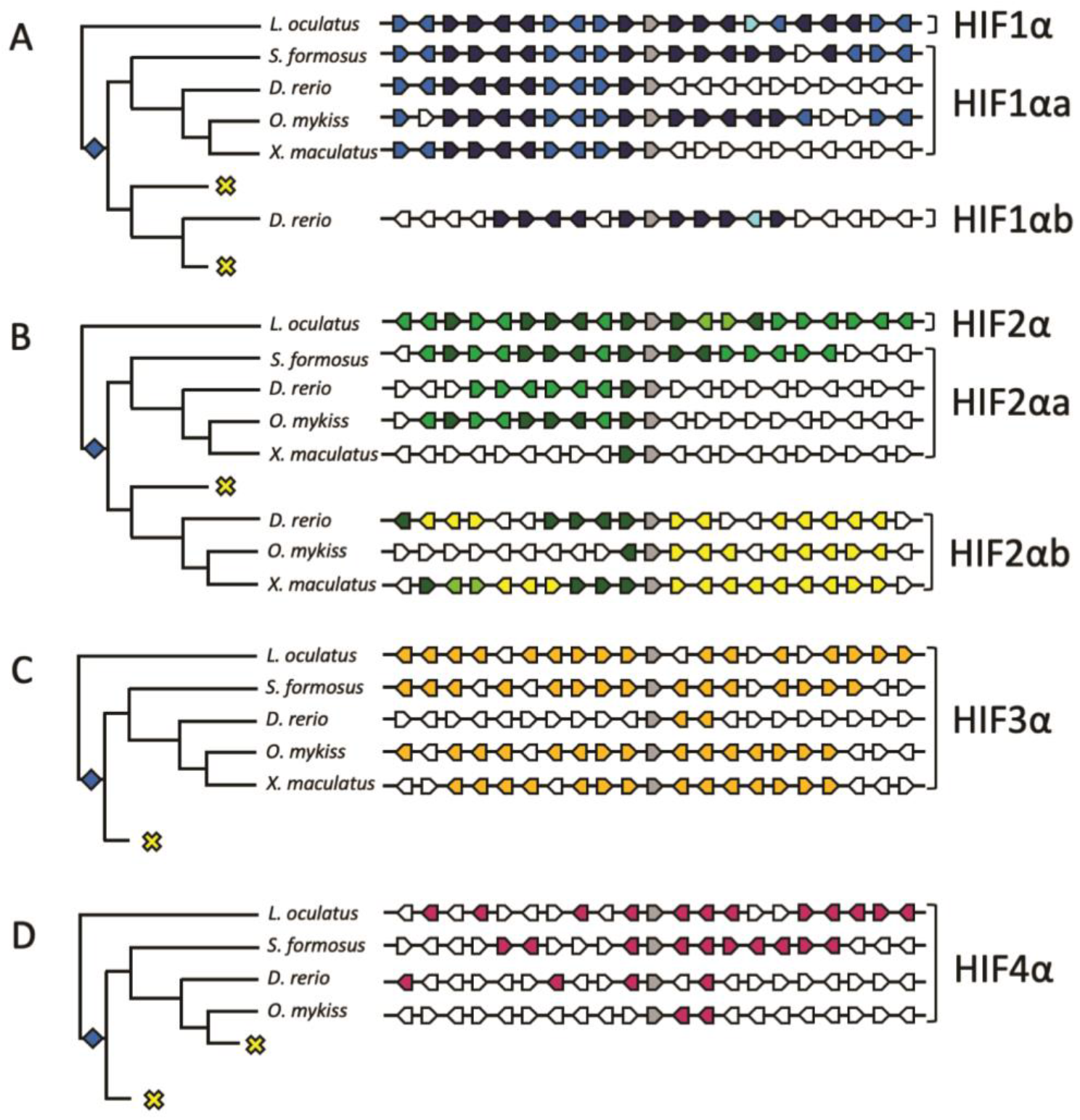
Synteny analysis of Actinopterygian HIF1α. The ten flanking genes on either side of each HIFα subunit (grey arrows) are compared among spotted gar (*L. oculatus*), Asian arowana (*S. formosus*), zebrafish (*D. rerio*), rainbow trout (*O. mykiss*), and southern platyfish (*X. maculatus*). A. Flanking genes shared between spotted gar HIF1α and both HIF1α paralogs in more derived fishes are shown as dark blue arrows, genes shared between spotted gar HIF1α and HIF1αa are shown as medium blue arrows, and genes shared between spotted gar HIF1α and HIF1αb are shown as light blue arrows. B. Flanking genes shared between spotted gar HIF2α and both HIF2α paralogs in more derived fishes are shown as dark green arrows, genes shared between spotted gar HIF2α and HIF2αa are shown as medium green arrows, and genes shared between spotted gar HIF2α and HIF2αb are shown as light green arrows. In addition, genes shared among HIF2αb, but not present in spotted gar, are shown as yellow arrows. For HIF3α (C) and HIF4α (D), flanking genes shared between spotted gar and more derived species are shown as orange and purple arrows, respectively. The relationships among fishes is from Hughes et al (2018) and branch lengths do not indicate divergence times. The teleost-specific genome duplication is indicated by the blue diamonds, and putative losses of specific HIFα forms are shown by the yellow crosses. For abbreviations of flanking genes, see supplementary table S2, Supplementary Material online.

Although the ancestral gene order flanking HIF2α is not as highly conserved across species, up to 10 genes are shared among one of the forms found in Otocephala and the truncated form found in more derived species (yellow arrows, fig. 2B). Rytkonen et al. (2013) originally proposed that this form be recognized as HIF2αb, a convention we previously endorsed (Townley et al. 2017) and maintain here (supplementary table S1, Supplementary Material online). Thus, the other paralog, which includes the first, full-length HIF2α described in fishes (Powell and Hahn 2002; Rojas et al. 2007), corresponds to HIF2αa. In current databases, however, the names of HIF2α paralogs are reversed: that is, the full-length and presumably functional form of HIF2α is listed as *hif2ab* or *epas1b*, whereas the other form, which is truncated in most fishes, is *hif2aa* or *epas1a*. Until there is consensus, great care will be needed when interpreting reports of paralog-specific differences in HIF2α.

Mining of fish genomes yielded only one form of HIF3α and HIF4α, and the latter was completely missing in more derived fishes (see above). The most parsimonious evolutionary scenario is that one paralog of each gene was lost shortly after the TGD, with the remaining form of HIF4α subsequently lost in Neoteleost (fig. 2C, D). An alternative hypothesis is that different teleost lineages retained different paralogs after the TGD (i.e., divergent silencing; Lynch 2002). This possibility is supported by the strong conservation of gene order flanking HIF3α within Otocephala, with very little shared synteny between Otocephala and other teleost lineages (fig 2C; supplemental table S2, Supplementary Material online). Further analyses will be required to confidently assign orthology of HIF3α and HIF4α in extant fishes.

As mentioned above, Salmoniformes have duplicates of HIF1α, HIF2α, and HIF3α arising from the SGD. We tentatively group these as “s1” and “s2” based upon sequence similarity within Salmoniformes and the observation that one of these (s1) shows greater shared synteny with the corresponding gene in Esociformes, the sister group of Salmoniformes (supplementary table S2, Supplementary Material online).

Among all species examined, the number of exons and length of deduced proteins are highly conserved within each HIFα type, with the notable exception of the truncated form, HIF2αb (fig. 3A, B; supplementary table S3 Supplementary Material online). We also analyzed variation in the amino acid sequences of putative functional domains, including the N-terminal and C-terminal oxygen dependent degradation domains (NODD and CODD) that are potential targets of regulation by prolyl hydroxylases (Epstein et al. 2001) and the extreme C-terminal CEVN motif targeted by asparaginyl hydroxylase (Lando et al. 2002) (fig. 3C). In general, the CODD is more highly conserved than the NODD, which is absent in HIF4α. These observations support suggestions that the CODD is more critical in determining the oxygen-dependence of HIFα subunit degradation (Epstein et al. 2001; Zhang et al. 2014). Within the CODD, however, sequence variation among forms of HIFα may be a source of functional divergence among ohnologs. For example, M_n-3_ is found in HIF1α and HIF3α and G_n+6_ is found in HIF2α and HIF4α (where n is the P that is putatively hydroxylated), and both residues are important in mediating HIFα degradation (Tarade et al. 2019). Finally, the lack of the CEVN sequence in HIF3α (Gu et al. 1998; Duan 2016) and lack of the critical asparagine in salmonid HIF4α suggests that these subunits are not subject to regulation by asparaginyl hydroxylation.

**Fig. 3.**
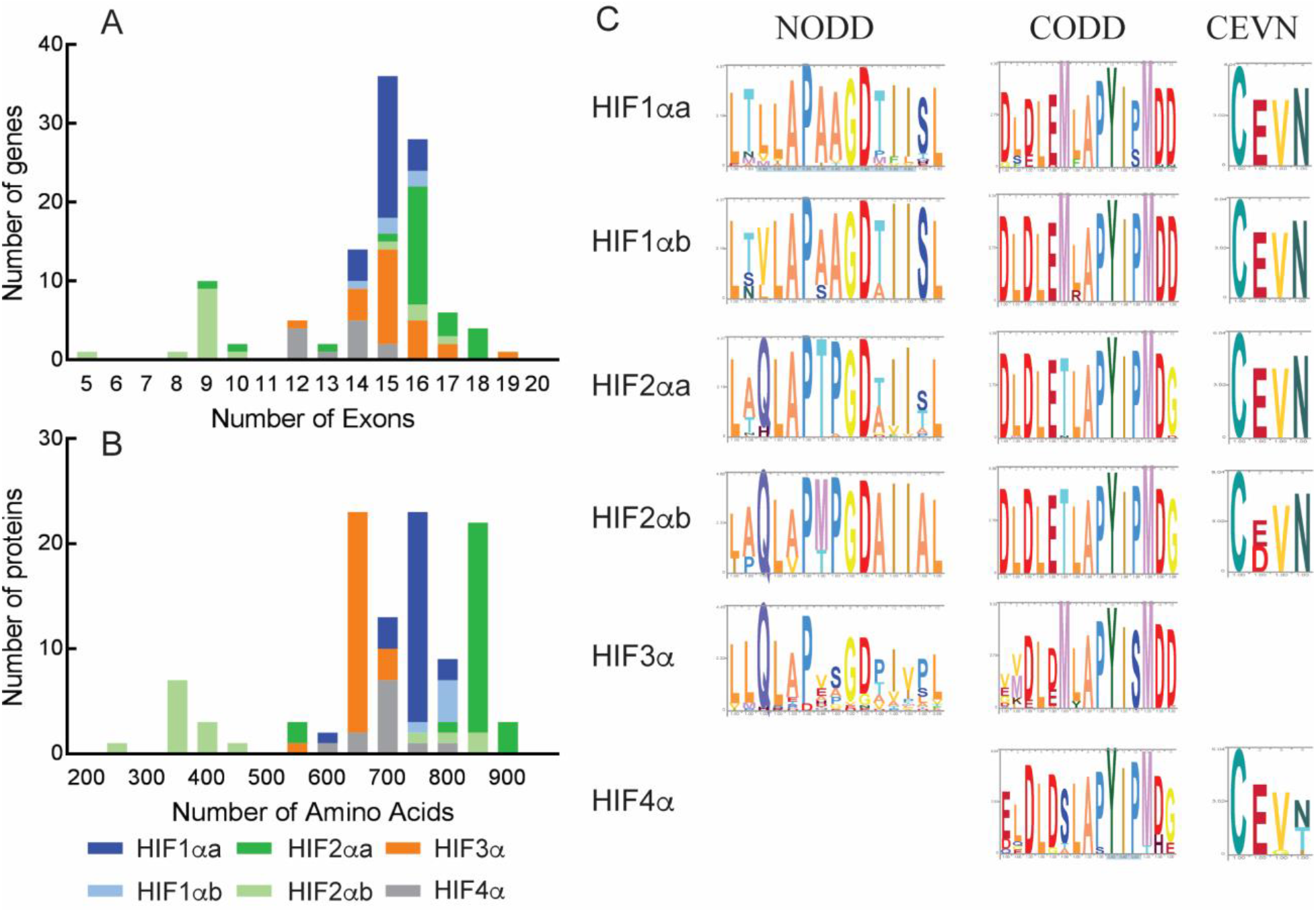
Exon number (A), deduced protein length (B), and consensus sequences for putative hydroxylation domains (C) for teleost HIFα subunits. Median values for exon number and amino acid number were as follows: HIF1αa = 15 exons, 759 amino acids (n=26); HIF1αb, 15 exons, 777 amino acids (n=5); HIF2αa, 16 exons, 853 amino acids (n=26); HIF3α, 15 exons, 646 amino acids (n=25); HIF4α, 14 exons, 716 amino acids (n=12). For HIF2αb, the distributions were bimodal, with median values of 16 exons and 816 amino acids for Otocephala (n=5) and 9 exons and 370 amino acids for other teleost (n=11). Exon number and deduced protein lengths for individual sequences are shown in supplementary table S1, Supplementary Material online, and summary statistics for major taxonomic groups are shown in supplementary table S3, Supplementary Material online. Consensus sequences for the N-and C-terminal oxygen-dependent degradation domains (NODD and CODD) and the C-terminal asparaginyl hydroxylation motif (CEVN) were identified from multiple sequence alignments and visualized with Skylign. The absence of a consensus sequence indicates that domain was not conserved for the respective HIFα subunit. Data for spotted gar (*Lepisosteus oculatus*) are not included.

## Conclusions

Here, we provide evidence that basal Actinopterygians retain four copies of HIFα that arose from the two rounds of GD at the base of vertebrate evolution. We advocate that a growing number of “HIFα-like” genes be recognized as HIF3α or HIF4α, as deduced by sequence analysis and shared synteny. The TGD and SGD in more derived fishes, followed by gene truncation and losses, resulted in the current diversity of HIFα among ray-finned fishes. Future research may better define the phylogenetic distribution and timing of gene loss events and determine whether different fish lineages retained different paralogs of HIF3α and HIF4α. In addition, functional analyses of HIFα paralogs, including truncated forms of HIF2αb, could illuminate the contribution of these duplicates to the oxygen relations and the general biology of ray-finned fishes. Hypotheses that the presence of gene duplicates confers enhanced tolerance of hypoxia, for example in cyprinids, should be re-examined in light of gene duplicates in closely related taxa (e.g., herring) or separate lineages (e.g., Salmoniformes) that are not particularly tolerant of low oxygen. It is plausible that these transcription factor duplicates have taken on functions apart from oxygen regulated gene expression (neofunctionalization) that may have contributed to the evolutionary and ecological success of Actinopterygii.

## Acknowledgements

We thank Jenna D. Hill and Nicholas J. Vogler for assistance in the initial stages of database searching and conserved domain analysis, respectively. We thank Drs. James Grady (University of New Orleans), Mark Hahn, Sibel Karchner (Woods Hole Oceanographic Institution), and Li Zhao (Rockefeller University) for constructive comments on this manuscript. This work was supported by the Greater New Orleans Foundation.

## Data Availability Statement

The data underlying this article are available at the National Center for Biotechnology Information at https://www.ncbi.nlm.nih.gov/ or at Ensembl at https://ensembl.org/index.html. All gene identification, mRNA, and protein accession numbers are provided in supplementary table S1, Supplementary Material online.

### Box 1. HIF gene symbols

Symbols for the hypoxia inducible factor alpha subunit follow the format:

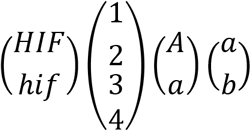

Generally, the following conventions apply:

1. The Human Genome Consortium Gene Nomenclature Committee specifies that gene symbols be all capital letters. The Zebrafish Nomenclature Conventions specifies lowercase Latin letters for gene symbols. For other species, uppercase, lowercase, or both have been used. The current study follows the convention for humans which has been broadly applied to other vertebrates and invertebrates (e.g., Graham and Presnell, 2017).
2. Paralogs that arose during two rounds of genome duplication at the base of vertebrate evolution are indicated by Arabic numbers (1-4).
3. Because electronic databases are not uniformly compatible with Greek letters, Latin letters (upper case in humans, lowercase in zebrafish) are used to distinguish the α subunit from the β subunit, which was originally described as the aryl hydrocarbon receptor nuclear translocator.
4. Paralogs that arose from the teleost-specific genome duplication are indicated by lowercase Latin letters (*a* or *b*).
5. Thus, the gene encoding the α subunit of the hypoxia inducible factor 1 is *HIF1A* in humans, but the two paralogs in zebrafish that arose from the TGD are *hif1aa* and *hif1ab*. Also, because the α subunit of the hypoxia inducible factor 2 was initially described as endothelial PAS-domain protein, this gene is also known as *EPAS1* in humans and *epas1a* or *epas1b* in zebrafish.

## References

Berthelot C, Brunet F, Chalopin D, Juanchich A, Bernard M, Noël B, Bento P, Da Silva C, Labadie K, Alberti A, et al. 2014. The rainbow trout genome provides novel insights into evolution after whole-genome duplication in vertebrates. Nat Commun. 5:3657. doi:10.1038/ncomms4657.

Bian C, Hu Y, Ravi V, Kuznetsova IS, Shen X, Mu X, Sun Y, You X, Li J, Li X, et al. 2016. The Asian arowana (Scleropages formosus) genome provides new insights into the evolution of an early lineage of teleosts. Sci Rep. 6:24501. doi:10.1038/srep24501.

Bouckaert R, Heled J, Kühnert D, Vaughan T, Wu C-H, Xie D, Suchard MA, Rambaut A, Drummond AJ. 2014. BEAST 2: A software platform for bayesian evolutionary analysis. PLOS Comput Biol. 10(4):e1003537. doi:10.1371/journal.pcbi.1003537.

Braasch I, Gehrke AR, Smith JJ, Kawasaki K, Manousaki T, Pasquier J, Amores A, Desvignes T, Batzel P, Catchen J, et al. 2016. The spotted gar genome illuminates vertebrate evolution and facilitates human-teleost comparisons. Nat Genet. 48(4):427–437. doi:10.1038/ng.3526.

Dehal P, Boore JL. 2005. Two rounds of whole genome duplication in the ancestral vertebrate. PLOS Biol. 3(10):e314. doi:10.1371/journal.pbio.0030314.

Drummond AJ, Ho SYW, Phillips MJ, Rambaut A. 2006. Relaxed Phylogenetics and Dating with Confidence. PLOS Biol. 4(5):e88. doi:10.1371/journal.pbio.0040088.

Drummond AJ, Suchard MA, Xie D, Rambaut A. 2012. Bayesian Phylogenetics with BEAUti and the BEAST 1.7. Mol Biol Evol. 29(8):1969–1973. doi:10.1093/molbev/mss075.

Duan C. 2016. Hypoxia-inducible factor 3 biology: complexities and emerging themes. Am J Physiol Cell Physiol. 310(4):C260–269. doi:10.1152/ajpcell.00315.2015.

Epstein AC, Gleadle JM, McNeill LA, Hewitson KS, O’Rourke J, Mole DR, Mukherji M, Metzen E, Wilson MI, Dhanda A, et al. 2001. C. elegans EGL-9 and mammalian homologs define a family of dioxygenases that regulate HIF by prolyl hydroxylation. Cell. 107(1):43–54. doi:10.1016/s0092-8674(01)00507-4.

Felsenstein J. 1985. Confidence Limits on Phylogenies: An Approach Using the Bootstrap. Evolution. 39(4):783–791. doi:10.1111/j.1558-5646.1985.tb00420.x.

Force A, Lynch M, Pickett FB, Amores A, Yan YL, Postlethwait J. 1999. Preservation of duplicate genes by complementary, degenerative mutations. Genetics. 151(4):1531–1545.

Glasauer SMK, Neuhauss SCF. 2014. Whole-genome duplication in teleost fishes and its evolutionary consequences. Mol Genet Genomics MGG. 289(6):1045–1060. doi:10.1007/s00438-014-0889-2.

Graham AM, Presnell JS. 2017. Hypoxia Inducible Factor (HIF) transcription factor family expansion, diversification, divergence and selection in eukaryotes. PloS One. 12(6):e0179545. doi:10.1371/journal.pone.0179545.

Gu YZ, Hogenesch JB, Bradfield CA. 2000. The PAS superfamily: sensors of environmental and developmental signals. Annu Rev Pharmacol Toxicol. 40:519–561. doi:10.1146/annurev.pharmtox.40.1.519.

Gu Y-Z, Moran SM, Hogenesch JB, Wartman L, Bradfield CA. 1998. Molecular characterization and chromosomal localization of a third α-class hypoxia inducible factor subunit, HIF3α. Gene Expr. 7(3):205–213.

Guo B. 2017. Complex genes are preferentially retained after whole-genome duplication in teleost fish. J Mol Evol. 84(5–6):253–258. doi:10.1007/s00239-017-9794-8.

Hughes LC, Ortí G, Huang Y, Sun Y, Baldwin CC, Thompson AW, Arcila D, Betancur-R R, Li C, Becker L, et al. 2018. Comprehensive phylogeny of ray-finned fishes (Actinopterygii) based on transcriptomic and genomic data. Proc Natl Acad Sci U S A. 115(24):6249–6254. doi:10.1073/pnas.1719358115.

Inoue J, Sato Y, Sinclair R, Tsukamoto K, Nishida M. 2015. Rapid genome reshaping by multiple-gene loss after whole-genome duplication in teleost fish suggested by mathematical modeling. Proc Natl Acad Sci U S A. 112(48):14918–14923. doi:10.1073/pnas.1507669112.

Jones DT, Taylor WR, Thornton JM. 1992. The rapid generation of mutation data matrices from protein sequences. Bioinformatics. 8(3):275–282. doi:10.1093/bioinformatics/8.3.275.

Kaelin WG, Ratcliffe PJ. 2008. Oxygen sensing by metazoans: the central role of the HIF hydroxylase pathway. Mol Cell. 30(4):393–402. doi:10.1016/j.molcel.2008.04.009.

Katoh K, Standley DM. 2013. MAFFT Multiple Sequence Alignment Software Version 7: Improvements in Performance and Usability. Mol Biol Evol. 30(4):772–780. doi:10.1093/molbev/mst010.

Katoh K, Standley DM. 2016. A simple method to control over-alignment in the MAFFT multiple sequence alignment program. Bioinformatics. 32(13):1933–1942. doi:10.1093/bioinformatics/btw108.

Kumar S, Stecher G, Li M, Knyaz C, Tamura K. 2018. MEGA X: Molecular Evolutionary Genetics Analysis across Computing Platforms. Mol Biol Evol. 35(6):1547–1549. doi:10.1093/molbev/msy096.

Lando D, Peet DJ, Gorman JJ, Whelan DA, Whitelaw ML, Bruick RK. 2002. FIH-1 is an asparaginyl hydroxylase enzyme that regulates the transcriptional activity of hypoxia-inducible factor. Genes Dev. 16(12):1466–1471. doi:10.1101/gad.991402.

Lynch M. 2002. Gene duplication and evolution. Science. 297(5583):945–947. doi:10.1126/science.1075472.

Lynch M, Conery JS. 2000. The evolutionary fate and consequences of duplicate genes. Science. 290(5494):1151–1155. doi:10.1126/science.290.5494.1151.

Macqueen DJ, Johnston IA. 2014. A well-constrained estimate for the timing of the salmonid whole genome duplication reveals major decoupling from species diversification. Proc R Soc B Biol Sci. 281(1778):20132881. doi:10.1098/rspb.2013.2881.

Makino Y, Kanopka A, Wilson WJ, Tanaka H, Poellinger L. 2002. Inhibitory PAS domain protein (IPAS) is a hypoxia-inducible splicing variant of the hypoxia-inducible factor-3alpha locus. J Biol Chem. 277(36):32405–32408. doi:10.1074/jbc.C200328200.

McIntosh BE, Hogenesch JB, Bradfield CA. 2010. Mammalian Per-Arnt-Sim proteins in environmental adaptation. Annu Rev Physiol. 72(1):625–645. doi:10.1146/annurev-physiol-021909-135922.

Nikinmaa M, Rees BB. 2005. Oxygen-dependent gene expression in fishes. Am J Physiol Regul Integr Comp Physiol. 288(5):R1079–1090. doi:10.1152/ajpregu.00626.2004.

Ohno S. 1970. Evolution by gene duplication. Berlin Heidelberg: Springer-Verlag.

Parey, E, Louis, A, Cabau, C, Gulgenn, Y, Crollius, HR, Berthelot, C. 2020. Synteny-guided resolution of gene trees clarifies the functional impact of whole genome duplications. Mol Biol Evol, doi.org/10.1093/molbev/msaa149

Pasquier J, Cabau C, Nguyen T, Jouanno E, Severac D, Braasch I, Journot L, Pontarotti P, Klopp C, Postlethwait JH, et al. 2016. Gene evolution and gene expression after whole genome duplication in fish: the PhyloFish database. BMC Genomics. 17:368. doi:10.1186/s12864-016-2709-z.

Penn O, Privman E, Ashkenazy H, Landan G, Graur D, Pupko T. 2010. GUIDANCE: a web server for assessing alignment confidence scores. Nucleic Acids Res. 38(suppl_2):W23–W28. doi:10.1093/nar/gkq443.

Postlethwait JH, Woods IG, Ngo-Hazelett P, Yan YL, Kelly PD, Chu F, Huang H, Hill-Force A, Talbot WS. 2000. Zebrafish comparative genomics and the origins of vertebrate chromosomes. Genome Res. 10(12):1890–1902. doi:10.1101/gr.164800.

Powell WH, Hahn ME. 2002. Identification and functional characterization of hypoxia-inducible factor 2alpha from the estuarine teleost, Fundulus heteroclitus: interaction of HIF-2alpha with two ARNT2 splice variants. J Exp Zool. 294(1):17–29. doi:10.1002/jez.10074.

Ravi V, Venkatesh B. 2018. The divergent genomes of teleosts. Annu Rev Anim Biosci. 6:47–68. doi:10.1146/annurev-animal-030117-014821.

Rojas DA, Perez-Munizaga DA, Centanin L, Antonelli M, Wappner P, Allende ML, Reyes AE. 2007. Cloning of hif-1alpha and hif-2alpha and mRNA expression pattern during development in zebrafish. Gene Expr Patterns GEP. 7(3):339–345. doi:10.1016/j.modgep.2006.08.002.

Rytkönen KT, Akbarzadeh A, Miandare HK, Kamei H, Duan C, Leder EH, Williams TA, Nikinmaa M. 2013. Subfunctionalization of cyprinid hypoxia-inducible factors for roles in development and oxygen sensing. Evol Int J Org Evol. 67(3):873–882. doi:10.1111/j.1558-5646.2012.01820.x.

Rytkönen KT, Williams TA, Renshaw GM, Primmer CR, Nikinmaa M. 2011. Molecular evolution of the metazoan PHD-HIF oxygen-sensing system. Mol Biol Evol. 28(6):1913–1926. doi:10.1093/molbev/msr012.

Sacerdot C, Louis A, Bon C, Berthelot C, Roest Crollius H. 2018. Chromosome evolution at the origin of the ancestral vertebrate genome. Genome Biol. 19(1):166. doi:10.1186/s13059-018-1559-1.

Sela I, Ashkenazy H, Katoh K, Pupko T. 2015. GUIDANCE2: accurate detection of unreliable alignment regions accounting for the uncertainty of multiple parameters. Nucleic Acids Res. 43(W1):W7–W14. doi:10.1093/nar/gkv318.

Semenza GL. 2011. Oxygen sensing, homeostasis, and disease. N Engl J Med. 365(6):537–547. doi:10.1056/NEJMra1011165.

Tarade D, Lee JE, Ohh M. 2019. Evolution of metazoan oxygen-sensing involved a conserved divergence of VHL affinity for HIF1α and HIF2α. Nature Comm 10:3293. doi.org/10.1038/s41467-019-11149-1

Townley IK, Karchner SI, Skripnikova E, Wiese TE, Hahn ME, Rees BB. 2017. Sequence and functional characterization of hypoxia-inducible factors, HIF1α, HIF2αa, and HIF3α, from the estuarine fish, Fundulus heteroclitus. Am J Physiol Regul Integr Comp Physiol. 312(3):R412–R425. doi:10.1152/ajpregu.00402.2016.

Volff J-N. 2005. Genome evolution and biodiversity in teleost fish. Heredity. 94(3):280–294. doi:10.1038/sj.hdy.6800635.

Waterhouse AM, Procter JB, Martin DMA, Clamp M, Barton GJ. 2009. Jalview Version 2--a multiple sequence alignment editor and analysis workbench. Bioinforma Oxf Engl. 25(9):1189–1191. doi:10.1093/bioinformatics/btp033.

Wheeler TJ, Clements J, Finn RD. 2014. Skylign: a tool for creating informative, interactive logos representing sequence alignments and profile hidden Markov models. BMC Bioinformatics. 15(1):7. doi:10.1186/1471-2105-15-7.

Wolfe, K. 2000. Robustness--it’s not where you think it is. Nat Genet 25: 3–4. doi.org/10.1038/75560

Zhang P, Yao Q, Lu L, Li Y, Chen P-J, Duan C. 2014. Hypoxia-inducible factor 3 is an oxygen-dependent transcription activator and regulates a distinct transcriptional response to hypoxia. Cell Rep. 6(6):1110–1121. doi:10.1016/j.celrep.2014.02.011.

