## Supplemental Figures for "Genomic Analysis of Hypoxia Inducible Factor Alpha Evolution in Ray-finned Fishes (Actinopterygii)"

**Fig. S1. Phylogeny of Actinopterygian HIFα reconstructed by Bayesian inference using full-length CDS.** Evolutionary analyses were conducted in BEAST 2 (v2.6.1) using the General Time Reversible model (GTR) with 6 gamma categories (+G) and allowing for invariants (+I). The tree with maximum clade credibility and mean heights is shown with posterior probability values next to the branches. The highest tree likelihood was -41437.47 with an ESS of 587. The tree was re-rooted on the outgroup for visualization. Four HIFα homology groups are indicated to the right and the following taxa are color coded within each group: Otocephala (green); Salmoniformes (orange); Neoteleostei (blue). The outgroup, *Ciona intestinalis*, basal Actinopterygian (spotted gar, *Lepisosteus oculatus*), basal teleost (Asian arowana, *Scleropages formosus*), and sister taxa to Salmoniformes (Northern pike, *Esox lucius*), are not color coded. Sequences are identified by the first letter of the genus and species followed by the last four digits of the NCBI or Ensemble reference gene accession number (see supplementary table S1, Supplementary Material online for a full list of genes). **NOTE:** this is a rectangular representation of the tree as shown in fig. 1 in circular format.

**Fig. S2. Phylogeny of Actinopterygian HIFα reconstructed by maximum likelihood using full-length CDS.** Evolutionary analyses were conducted in MEGAX (v10.1.8) using the General Time Reversible model (GTR) with 6 gamma categories(+G) and allowing for invariants (+I). The tree with the highest log likelihood (-41489.7) is shown with bootstrap values next to the branches. The tree was re-rooted on the outgroup for visualization. Four HIFα homology groups are indicated to the right and the following taxa are color coded within each group: Otocephala (green); Salmoniformes (orange); Neoteleostei (blue). The outgroup, *Ciona intestinalis*, basal Actinopterygian (spotted gar, *Lepisosteus oculatus*), basal teleost (Asian arowana, *Scleropages formosus*), and sister taxa to Salmoniformes (Northern pike, *Esox lucius*), are not color coded. Sequences are identified by the first letter of the genus and species followed by the last four digits of the NCBI or Ensemble reference gene accession number (see supplementary table S1, Supplementary Material online for a full list of genes).

**Fig. S3. Phylogeny of Actinopterygian HIFα reconstructed by Bayesian inference using amino acid sequences deduced for full length proteins.** Evolutionary analyses were conducted in BEAST 2 (v2.6.1) using a JTT matrix-based model with 6 gamma categories (+G) and allowing for invariants (+I). The tree with maximum clade credibility and mean heights is shown with posterior probability values next to the branches. The highest tree likelihood was -20378.55 with an ESS of 964.The tree was re-rooted on the outgroup for visualization. Four HIFα homology groups are indicated to the right and the following taxa are color coded within each group: Otocephala (green); Salmoniformes (orange); Neoteleostei (blue). The outgroup, *Ciona intestinalis*, basal Actinopterygian (spotted gar, *Lepisosteus oculatus*), basal teleost (Asian arowana, *Scleropages formosus*), and sister taxa to Salmoniformes (Northern pike, *Esox lucius*), are not color coded. Sequences are identified by the first letter of the genus and species followed by the last four digits of the NCBI or Ensemble reference gene accession number (see supplementary table S1, Supplementary Material online for a full list of genes).

**Fig. S4. Phylogeny of Actinopterygian HIFα reconstructed by maximum likelihood using amino acid sequences deduced for full length proteins.** Evolutionary analyses were conducted in MEGAX (v10.1.8) using a JTT matrix-based model with 6 gamma categories (+G) and allowing for invariants (+I). The tree with the highest log likelihood (-20259.51) is shown with bootstrap values next to the branches. The tree was re-rooted on the outgroup after analysis. Four HIFα homology groups are indicated to the right and the following taxa are color coded within each group: Otocephala (green); Salmoniformes (orange); Neoteleostei (blue). The outgroup, *Ciona intestinalis*, basal Actinopterygian (spotted gar, *Lepisosteus oculatus*), basal teleost (Asian arowana, *Scleropages formosus*), and sister taxa to Salmoniformes (Northern pike, *Esox lucius*), are not color coded. Sequences are identified by the first letter of the genus and species followed by the last four digits of the NCBI or Ensemble reference gene accession number (see supplementary table S1, Supplementary Material online for a full list of genes).

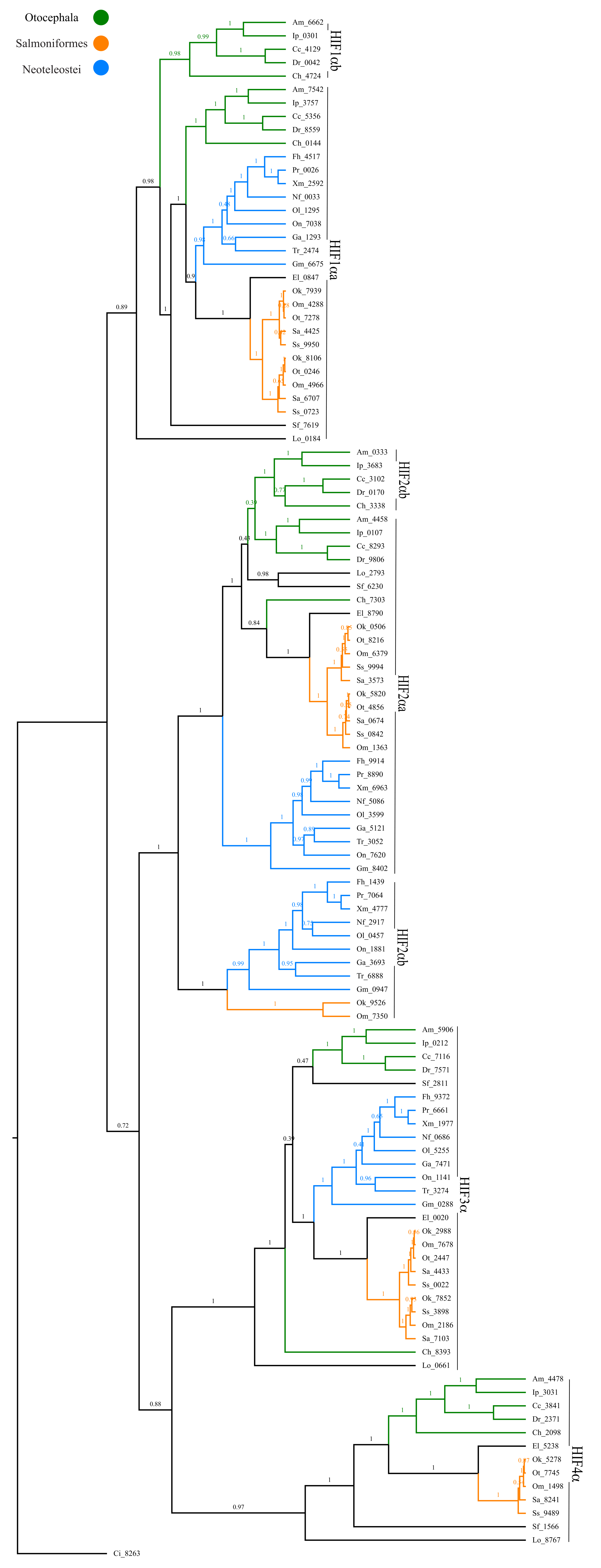
Otocephala
Salmoniformes
Neoteleostei
HIF1αb
HIF1αa
HIF2αb
HIF2αa
HIF2αb
HIF3α
HIF4α
Ci_8263

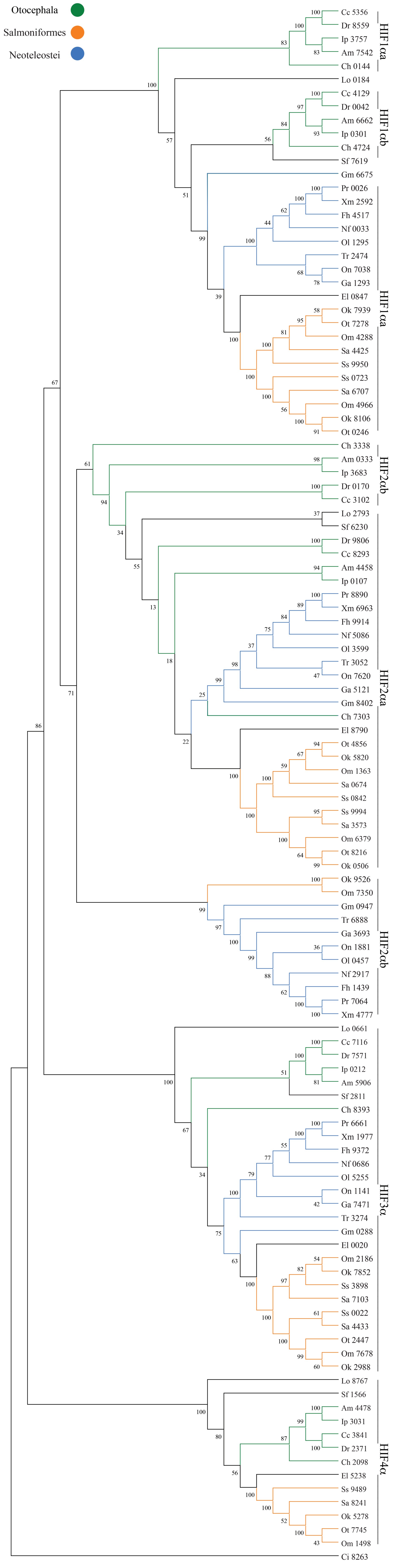
Otocephala
Salmoniformes
Neoteleostei
Cc 5356
Dr 8559
Ip 3757
Am 7542
Ch 0144
HIF1αa
Lo 0184
Cc 4129
Dr 0042
Am 6662
Ip 0301
Ch 4724
Sf 7619
HIF1αb
Gm 6675
Pr 0026
Xm 2592
Fh 4517
Nf 0033
Ol 1295
Tr 2474
On 7038
Ga 1293
El 0847
Ok 7939
Ot 7278
Om 4288
Sa 4425
Ss 9950
Ss 0723
Sa 6707
Om 4966
Ok 8106
Ot 0246
HIF1αa
Ch 3338
Am 0333
Ip 3683
Dr 0170
Cc 3102
HIF2αb
Lo 2793
Sf 6230
Dr 9806
Cc 8293
Am 4458
Ip 0107
Pr 8890
Xm 6963
Fh 9914
Nf 5086
Ol 3599
Tr 3052
On 7620
Ga 5121
Gm 8402
Ch 7303
El 8790
Ot 4856
Ok 5820
Om 1363
Sa 0674
Ss 0842
Ss 9994
Sa 3573
Om 6379
Ot 8216
Ok 0506
HIF2αa
Ok 9526
Om 7350
Gm 0947
Tr 6888
Ga 3693
On 1881
Ol 0457
Nf 2917
Fh 1439
Pr 7064
Xm 4777
HIF2αb
Lo 0661
Cc 7116
Dr 7571
Ip 0212
Am 5906
Sf 2811
Ch 8393
Pr 6661
Xm 1977
Fh 9372
Nf 0686
Ol 5255
On 1141
Ga 7471
Tr 3274
Gm 0288
El 0020
Om 2186
Ok 7852
Ss 3898
Sa 7103
Ss 0022
Sa 4433
Ot 2447
Om 7678
Ok 2988
HIF3α
Lo 8767
Sf 1566
Am 4478
Ip 3031
Cc 3841
Dr 2371
Ch 2098
El 5238
Ss 9489
Sa 8241
Ok 5278
Ot 7745
Om 1498
HIF4α
Ci 8263

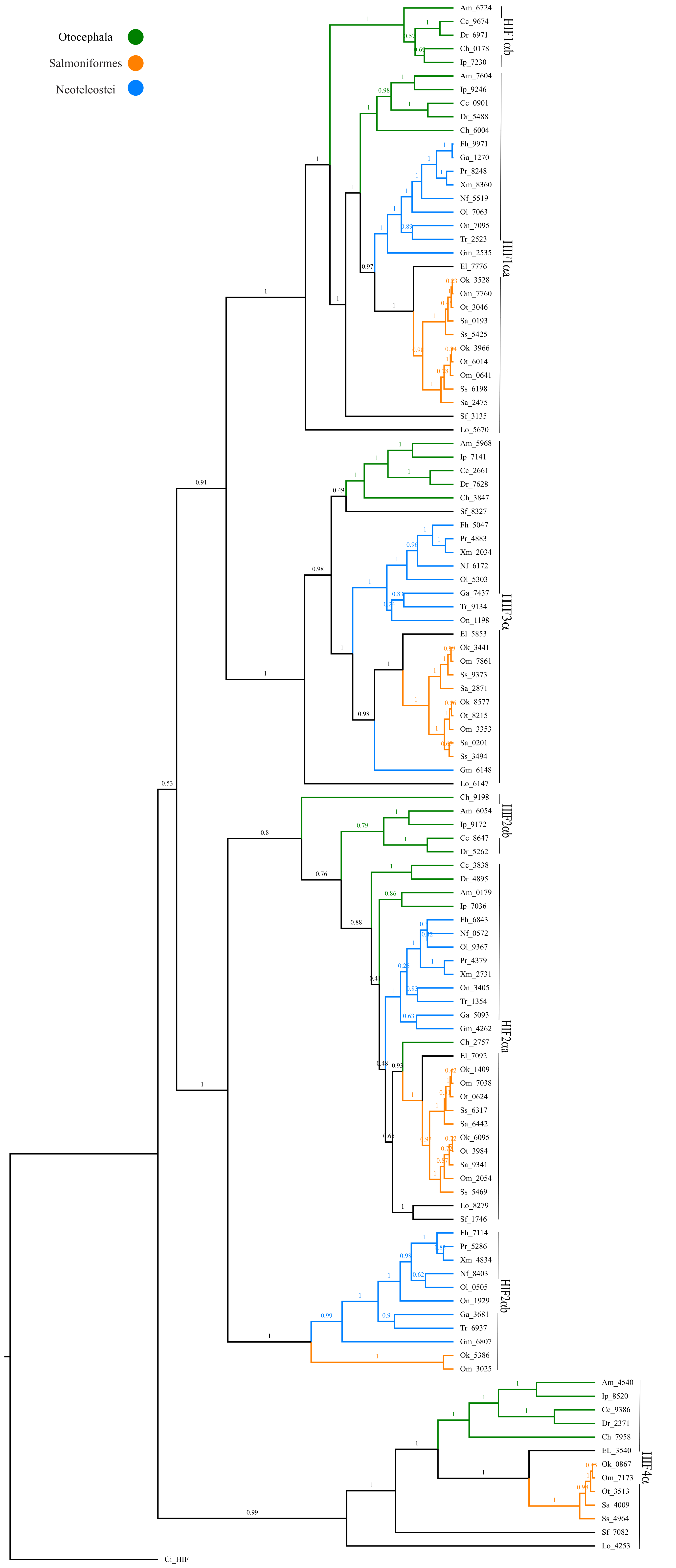
Otocephala
Salmoniformes
Neoteleostei
Am_6724
Cc_9674
Dr_6971
Ch_0178
Ip_7230
HIF1αb
Am_7604
Ip_9246
Cc_0901
Dr_5488
Ch_6004
Fh_9971
Ga_1270
Pr_8248
Xm_8360
Nf_5519
Ol_7063
On_7095
Tr_2523
Gm_2535
El_7776
Ok_3528
Om_7760
Ot_3046
Sa_0193
Ss_5425
Ok_3966
Ot_6014
Om_0641
Ss_6198
Sa_2475
Sf_3135
Lo_5670
HIF1αa
Am_5968
Ip_7141
Cc_2661
Dr_7628
Ch_3847
Sf_8327
Fh_5047
Pr_4883
Xm_2034
Nf_6172
Ol_5303
Ga_7437
Tr_9134
On_1198
El_5853
Ok_3441
Om_7861
Ss_9373
Sa_2871
Ok_8577
Ot_8215
Om_3353
Sa_0201
Ss_3494
Gm_6148
Lo_6147
HIF3α
Ch_9198
Am_6054
Ip_9172
Cc_8647
Dr_5262
HIF2αb
Cc_3838
Dr_4895
Am_0179
Ip_7036
Fh_6843
Nf_0572
Ol_9367
Pr_4379
Xm_2731
On_3405
Tr_1354
Ga_5093
Gm_4262
Ch_2757
El_7092
Ok_1409
Om_7038
Ot_0624
Ss_6317
Sa_6442
Ok_6095
Ot_3984
Sa_9341
Om_2054
Ss_5469
Lo_8279
Sf_1746
HIF2αa
Fh_7114
Pr_5286
Xm_4834
Nf_8403
Ol_0505
On_1929
Ga_3681
Tr_6937
Gm_6807
Ok_5386
Om_3025
HIF2αb
Am_4540
Ip_8520
Cc_9386
Dr_2371
Ch_7958
EL_3540
Ok_0867
Om_7173
Ot_3513
Sa_4009
Ss_4964
Sf_7082
Lo_4253
HIF4α
Ci_HIF

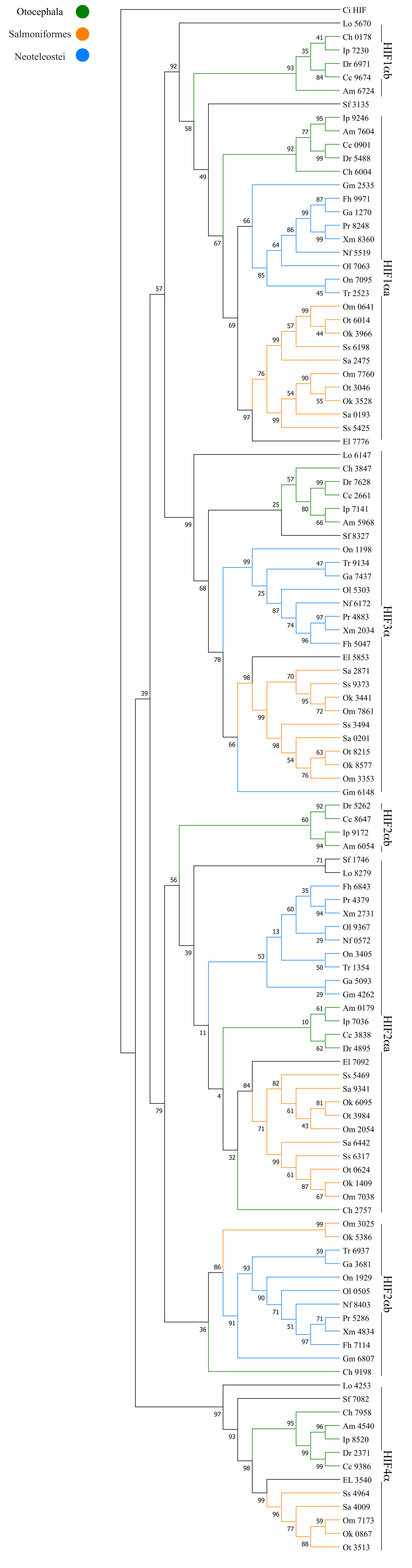
Otocephala
Salmoniformes
Neoteleostei
Ci HIF
Lo 5670
Ch 0178
Ip 7230
Dr 6971
Cc 9674
Am 6724
HIF1αb
Sf 3135
Ip 9246
Am 7604
Cc 0901
Dr 5488
Ch 6004
Gm 2535
Fh 9971
Ga 1270
Pr 8248
Xm 8360
Nf 5519
Ol 7063
On 7095
Tr 2523
Om 0641
Ot 6014
Ok 3966
Ss 6198
Sa 2475
Om 7760
Ot 3046
Ok 3528
Sa 0193
Ss 5425
El 7776
HIF1αa
Lo 6147
Ch 3847
Dr 7628
Cc 2661
Ip 7141
Am 5968
Sf 8327
On 1198
Tr 9134
Ga 7437
Ol 5303
Nf 6172
Pr 4883
Xm 2034
Fh 5047
El 5853
Sa 2871
Ss 9373
Ok 3441
Om 7861
Ss 3494
Sa 0201
Ot 8215
Ok 8577
Om 3353
Gm 6148
HIF3α
Dr 5262
Cc 8647
Ip 9172
Am 6054
HIF2αb
Sf 1746
Lo 8279
Fh 6843
Pr 4379
Xm 2731
Ol 9367
Nf 0572
On 3405
Tr 1354
Ga 5093
Gm 4262
Am 0179
Ip 7036
Cc 3838
Dr 4895
El 7092
Ss 5469
Sa 9341
Ok 6095
Ot 3984
Om 2054
Sa 6442
Ss 6317
Ot 0624
Ok 1409
Om 7038
Ch 2757
HIF2αa
Om 3025
Ok 5386
Tr 6937
Ga 3681
On 1929
Ol 0505
Nf 8403
Pr 5286
Xm 4834
Fh 7114
Gm 6807
Ch 9198
HIF2αb
Lo 4253
Sf 7082
Ch 7958
Am 4540
Ip 8520
Dr 2371
Cc 9386
EL 3540
Ss 4964
Sa 4009
Om 7173
Ok 0867
Ot 3513
HIF4α
