## Supplemental Tables for "Genomic Analysis of Hypoxia Inducible Factor Alpha Evolution in Ray-finned Fishes (Actinopterygii)"

**Supplemental Table Legends**

**Table S1. List of HIFα sequences used for phylogenetic analyses.** For each sequence, the organism name (species and common name) and order (*sensu* Hughes et al. 2018) are given. Gene name, aliases, Gene ID, mRNA accession, protein accession, and gene and protein statistics are from GenBank, except for three-spined stickleback (*Gasterosteus aculeatus*) which are from Ensemble. The gene group is based upon the phylogenetic and synteny analyses reported here (figure 1, electronic supplementary material, table S2). When multiple variants were identified within a locus, the longest form was selected. **NOTE:** data for spotted gar (*Lepisosteus oculatus*) HIF1α were generated by concatenation of two partial sequences (Gene ID 107077742 and 102694568).

**Table S2. Synteny analysis of Actinopterygian HIFα subunits.** The ten flanking genes on either side of each HIFα subunit were determined for spotted gar (*L. oculatus*), Asian arowana (*S. formosus*), representative Otocephala (*D. rerio* and *A. mexicanus*), northern pike (*E. lucius*), rainbow trout (*O. mykiss*), and representative Neoteleost (*X. maculatus* and *T. rubripes*). Flanking genes shared between spotted gar HIF1α and both HIF1α paralogs in more derived fishes are highlighted in dark blue, genes shared between spotted gar HIF1α and HIF1αa are highlighted in medium blue, and genes shared between spotted gar HIF1α and HIF1αb are highlighted in light blue. Flanking genes shared between spotted gar HIF2α and both HIF2α paralogs in more derived fishes are highlighted in dark green, genes shared between spotted gar HIF2α and HIF2αa are highlighted in medium green, and genes shared between spotted gar HIF2α and HIF2αb are highlighted in light green. In addition, genes shared among HIF2αb, but not present in spotted gar, are highlighted in yellow. Flanking genes shared between spotted gar HIF3α and HIF3α from more derived species are highlighted in orange. Flanking genes shared between spotted gar HIF4α and HIF4α from more derived species are highlighted in purple. Gene ID, chromosomal or linkage group assignment, mRNA accession, and protein accession are from GenBank. Gene orientation indicated by the forward or reverse arrow heads (-> or <-).

**Table S3. Exon number and deduced protein length of Actinopterygian HIFα subunits.** HIFα group is based upon phylogenetic analysis (figure 1) and synteny (figure 2, electronic supplementary material, table S2). Within each HIFα, species are grouped according to phylogeny (Hughes et al. 2018). The number of species in each group is indicated, along with the minimum, maximum, and median values for exon number and amino acid number for the deduced protein sequence. When multiple variants were identified within a locus, the longest form was selected. Data for spotted gar (*Lepisosteus oculatus*) are not included.

| Species | Common Name | Order | Genbank or Ensemble Gene Name | Group | Aliases | Gene ID (Genbank) | Chromosome # | Exons | mRNA Accession No. | Protein Accession No. | Protein Length (aa) |
|---|---|---|---|---|---|---|---|---|---|---|---|
| *Ciona intestinalis* | sea squirt | OUTGROUP | transcription factor protein | HIF | hif | 778640 | 4 | 22 | NM_001078263 | NP_001071731 | 735 |
| *Astyanax mexicanus* | cave fish | Characiformes | hypoxia inducible factor 1 subunit alpha a | HIF1Aa | hif1aa | 103022448 | 25 | 15 | XM_007247542 | XP_007247604 | 721 |
| *Astyanax mexicanus* | cave fish | Characiformes | hypoxia inducible factor 1 subunit alpha b | HIF1Ab | hif1ab | 103033873 | 3 | 14 | XM_007256662 | XP_007256724 | 785 |
| *Astyanax mexicanus* | cave fish | Characiformes | endothelial PAS domain protein 1b | HIF2Aa | epas1b | 103026643 | 25 | 17 | XM_022684458 | XP_022540179 | 912 |
| *Astyanax mexicanus* | cave fish | Characiformes | endothelial PAS domain protein 1a | HIF2Ab | epas1a | 103036701 | 6 | 16 | XM_022670333 | XP_022526054 | 816 |
| *Astyanax mexicanus* | cave fish | Characiformes | hypoxia inducible factor 1 subunit alpha, like | HIF3A | hif1al | 103027586 | 1 | 15 | XM_007245906 | XP_007245968 | 633 |
| *Astyanax mexicanus* | cave fish | Characiformes | hypoxia inducible factor 1 subunit alpha, like 2 | HIF4A | hif1al2 | 103041845 | 14 | 14 | XM_007244478 | XP_007244540 | 661 |
| *Cyprinus carpio* | common carp | Cypriniformes | hypoxia inducible factor 1 subunit alpha a | HIF1Aa | hif1aa | 109112424 | 38 | 15 | XM_019125356 | XP_018980901 | 712 |
| *Cyprinus carpio* | common carp | Cypriniformes | hypoxia-inducible factor 1-alpha-like | HIF1Ab | hif1al | 109100693 | 25 | 15 | XM_019114129 | XP_018969674 | 774 |
| *Cyprinus carpio* | common carp | Cypriniformes | endothelial PAS domain-containing protein 1-like | HIF2Aa | | 109094582 | 15 | 10 | XM_019108293 | XP_018963838 | 565 |
| *Cyprinus carpio* | common carp | Cypriniformes | endothelial PAS domain protein 1a | HIF2Ab | epas1a | 109099593 | 23 | 17 | XM_019113102 | XP_018968647 | 845 |
| *Cyprinus carpio* | common carp | Cypriniformes | hypoxia inducible factor 1 subunit alpha, like | HIF3A | hif1al | 109103773 | 30 | 17 | XM_019117116 | XP_018972661 | 635 |
| *Cyprinus carpio* | common carp | Cypriniformes | endothelial PAS domain-containing protein 1-like | HIF4A | | 109056647 | un | 14 | XM_019073841 | XP_018929386 | 676 |
| *Clupea harengus* | Atlantic herring | Clupeiformes | hypoxia-inducible factor 1-alpha-like | HIF1Aa | hif1al | 105901315 | 14 | 15 | XM_031580144 | XP_031436004 | 748 |
| *Clupea harengus* | Atlantic herring | Clupeiformes | hypoxia inducible factor 1 subunit alpha b | HIF1Ab | hif1ab | 105897766 | 15 | 16 | XM_012824724 | XP_012680178 | 798 |
| *Clupea harengus* | Atlantic herring | Clupeiformes | endothelial PAS domain protein 1b | HIF2Aa | epas1b | 105900051 | 13 | 16 | XM_012827303 | XP_012682757 | 838 |
| *Clupea harengus* | Atlantic herring | Clupeiformes | endothelial PAS domain-containing protein 1-like | HIF2Ab | | 105904118 | 16 | 10 | XM_031583338 | XP_031439198 | 459 |
| *Clupea harengus* | Atlantic herring | Clupeiformes | hypoxia inducible factor 1 subunit alpha, like | HIF3A | hif1al | 105909740 | 9 | 15 | XM_012838393 | XP_012693847 | 646 |
| *Clupea harengus* | Atlantic herring | Clupeiformes | hypoxia inducible factor 1 subunit alpha, like 2 | HIF4A | hif1al2 | 105900319 | 8 | 14 | XM_031572098 | XP_031427958 | 708 |
| *Danio rerio* | zebrafish | Cypriniformes | hypoxia inducible factor 1 subunit alpha a | HIF1Aa | hif1aa | 797150 | 13 | 15 | NM_001308559 | NP_001295488 | 717 |
| *Danio rerio* | zebrafish | Cypriniformes | hypoxia inducible factor 1 subunit alpha b | HIF1Ab | hif1ab | 393292 | 20 | 15 | NM_001310042 | NP_001296971 | 777 |
| *Danio rerio* | zebrafish | Cypriniformes | endothelial PAS domain protein 1b | HIF2Aa | epas1b | 555192 | 13 | 16 | NM_001039806 | NP_001034895 | 834 |
| *Danio rerio* | zebrafish | Cypriniformes | endothelial PAS domain protein 1a | HIF2Ab | epas1a | 566886 | 12 | 16 | XM_690170 | XP_695262 | 843 |
| *Danio rerio* | zebrafish | Cypriniformes | hypoxia inducible factor 1 subunit alpha, like | HIF3A | hif1al | 393376 | 15 | 17 | XM_005157571 | XP_005157628 | 626 |
| *Danio rerio* | zebrafish | Cypriniformes | hypoxia inducible factor 1 subunit alpha, like 2 | HIF4A | hif1al2 | 497283 | 21 | 15 | NM_001012371 | NP_001012371 | 663 |
| *Esox lucius* | Northern pike | Esosiformes | hypoxia inducible factor 1 subunit alpha a | HIF1Aa | hif1a, hif1aa | 105015558 | 15 | 15 | NM_001310847 | NP_001297776 | 763 |
| *Esox lucius* | Northern pike | Esosiformes | endothelial PAS domain protein 1b | HIF2Aa | epas1, epas1b | 105021253 | 5 | 16 | XM_010888790 | XP_010887092 | 852 |
| *Esox lucius* | Northern pike | Esosiformes | hypoxia inducible factor 1 subunit alpha, like | HIF3A | hif1al | 105012786 | 1 | 15 | XM_029120020 | XP_028975853 | 649 |
| *Esox lucius* | Northern pike | Esosiformes | hypoxia inducible factor 1 subunit alpha, like 2 | HIF4A | hif1al2 | 105024933 | 7 | 14 | XM_010895238 | XP_010893540 | 775 |
| *Fundulus heteroclitius* | mummichog | Cyprinodontiformes | hypoxia-inducible factor 1 subunit alpha a | HIF1Aa | hif1aa | 105927646 | un | 15 | XM_012864517 | XP_012719971 | 754 |
| *Fundulus heteroclitius* | mummichog | Cyprinodontiformes | endothelial PAS domain protein 1b | HIF2Aa | epas1, epas1b | 105933321 | un | 16 | NM_001309914 | NP_001296843 | 873 |
| *Fundulus heteroclitius* | mummichog | Cyprinodontiformes | endothelial PAS domain protein 1a | HIF2Ab | epas1, epas1a | 105918888 | un | 9 | XM_021311439 | XP_021167114 | 370 |
| *Fundulus heteroclitius* | mummichog | Cyprinodontiformes | hypoxia inducible factor 1 subunit alpha, like | HIF3A | hif1al | 105930192 | un | 12 | XM_021319372 | XP_021175047 | 662 |
| *Gasterosteus aculeatus* | three-spined stickleback | Perciformes | hypoxia-inducible factor 1 alpha | HIF1Aa | hif-1a | ENSGACG00000008525 | LG15 | 16 | ENSGACT00000011293 | ENSGACP00000011270 | 760 |
| *Gasterosteus aculeatus* | three-spined stickleback | Perciformes | endeothelial PAS domain protein 1b | HIF2Aa | epas1b | ENSGACG00000011414 | LG6 | 18 | ENSGACT00000015121 | ENSGACP00000015093 | 834 |
| *Gasterosteus aculeatus* | three-spined stickleback | Perciformes | novel gene | HIF2Ab | | ENSGACG00000002809 | LG5 | 9 | ENSGACT00000003693 | ENSGACP00000003681 | 369 |
| *Gasterosteus aculeatus* | three-spined stickleback | Perciformes | hypoxia inducible factor 1 subunit alpha, like | HIF3A | hif1al | ENSGACG00000013185 | LG1 | 14 | ENSGACT00000017471 | ENSGACP00000017437 | 651 |
| *Gadus morhua* | Atlantic cod | Gadiformes | hypoxia inducible factor 1 subunit alpha a | HIF1Aa | hif1aa | 115543958 | 5 | 15 | XM_030356675 | XP_030212535 | 774 |
| *Gadus morhua* | Atlantic cod | Gadiformes | endothelial PAS domain protein 1b | HIF2Aa | epas1b | 115559556 | 15 | 16 | XM_030378402 | XP_030234262 | 841 |
| *Gadus morhua* | Atlantic cod | Gadiformes | endothelial PAS domain protein 1 | HIF2Ab | epas1 | 115531593 | 18 | 8 | XM_030340947 | XP_030196807 | 367 |
| *Gadus morhua* | Atlantic cod | Gadiformes | hypoxia inducible factor 1 subunit alpha, like | HIF3A | hif1al | 115560760 | 16 | 15 | XM_030380288 | XP_030236148 | 667 |
| *Ictalurus punctatus* | channel catfish | Siluriformes | hypoxia inducible factor 1 subunit alpha a | HIF1Aa | hif1aa | 108263197 | 3 | 14 | XM_017463757 | XP_017319246 | 611 |
| *Ictalurus punctatus* | channel catfish | Siluriformes | hypoxia inducible factor 1 subunit alpha b | HIF1Ab | hif1ab | 100305068 | 25 | 16 | NM_001200301 | NP_001187230 | 776 |
| *Ictalurus punctatus* | channel catfish | Siluriformes | endothelial PAS domain protein 1b | HIF2Aa | epas, epas1b | 100304992 | 3 | 16 | NM_001350107 | NP_001337036 | 820 |
| *Ictalurus punctatus* | channel catfish | Siluriformes | endothelial PAS domain protein 1a | HIF2Ab | epas1a | 108273968 | 13 | 15 | XM_017483683 | XP_017339172 | 760 |
| *Ictalurus punctatus* | channel catfish | Siluriformes | hypoxia inducible factor 1 subunit alpha, like | HIF3A | hif1al | 100304653 | 17 | 15 | NM_001200212 | NP_001187141 | 627 |
| *Ictalurus punctatus* | channel catfish | Siluriformes | hypoxia inducible factor 1 subunit alpha, like 2 | HIF4A | hif1al2 | 108279075 | 18 | 14 | XM_017493031 | XP_017348520 | 606 |
| *Lepisosteus oculatus* | spotted gar | Lepisosteiformes | hypoxia inducible factor 1 subunit alpha | HIF1A | hif1a | 102682513+107077742 | LG7 | 10+3 | XM_015350184+XM_015350128 | XP_015205670+XP_015205614 | 430+121 |
| *Lepisosteus oculatus* | spotted gar | Lepisosteiformes | endeothelial PAS domain protein 1b | HIF2A | epas1b | 102694568 | LG16 | 16 | XM_015362793 | XP_015218279 | 852 |
| *Lepisosteus oculatus* | spotted gar | Lepisosteiformes | hypoxia inducible factor 1 subunit alpha, like | HIF3A | hif1al | 102689100 | LG2 | 13 | XM_015340661 | XP_015196147 | 488 |
| *Lepisosteus oculatus* | spotted gar | Lepisosteiformes | endothelial PAS domain-containing protein 1-like | HIF4A | | 102687670 | LG28 | 9 | XM_015338767 | XP_015194253 | 527 |
| *Nothobranchius furzeri* | turquoise killifish | Cyprinodontiformes | hypoxia inducible factor 1 subunit alpha a | HIF1Aa | hif1aa | 107392302 | sgr16 | 16 | XM_015970033 | XP_015825519 | 748 |
| *Nothobranchius furzeri* | turquoise killifish | Cyprinodontiformes | endothelial PAS domain protein 1b | HIF2Aa | epas1b | 107376137 | sgr03 | 16 | XM_015945086 | XP_015800572 | 860 |
| *Nothobranchius furzeri* | turquoise killifish | Cyprinodontiformes | endothelial PAS domain protein 1a | HIF2Ab | epas1a | 107387754 | sgr12 | 9 | XM_015962917 | XP_015818403 | 364 |
| *Nothobranchius furzeri* | turquoise killifish | Cyprinodontiformes | hypoxia inducible factor 1 subunit alpha, like | HIF3A | hif1al | 107379773 | sgr06 | 15 | XM_015950686 | XP_015806172 | 649 |
| *Oncorhynchus kisutch* | coho salmon | Salmoniformes | hypoxia-inducible factor 1-alpha | HIF1Aa_s1 | | 109894443 | LG7 | 14 | XM_020487939 | XP_020343528 | 762 |
| *Oncorhynchus kisutch* | coho salmon | Salmoniformes | hypoxia inducible factor 1 subunit alpha a | HIF1Aa_s2 | hif1a | 109904113 | LG14 | 15 | XM_031788106 | XP_031643966 | 758 |
| *Oncorhynchus kisutch* | coho salmon | Salmoniformes | endothelial PAS domain protein 1 | HIF2Aa_s1 | epas1 | 109870143 | LG25 | 16 | XM_020460506 | XP_020316095 | 853 |
| *Oncorhynchus kisutch* | coho salmon | Salmoniformes | endothelial PAS domain-containing protein 1-like | HIF2Aa_s2 | | 109900095 | LG11 | 16 | XM_020495820 | XP_020351409 | 845 |
| *Oncorhynchus kisutch* | coho salmon | Salmoniformes | endothelial PAS domain-containing protein 1-like | HIF2Ab | | 109865102 | LG20 | 9 | XM_031799526 | XP_031655386 | 391 |
| *Oncorhynchus kisutch* | coho salmon | Salmoniformes | hypoxia-inducible factor 1-alpha-like | HIF3A_s1 | hif1al | 109905559 | LG15 | 16 | XM_020502988 | XP_020358577 | 627 |
| *Oncorhynchus kisutch* | coho salmon | Salmoniformes | hypoxia-inducible factor 3-alpha | HIF3A_s2 | | 109909008 | LG18 | 15 | XM_020507852 | XP_020363441 | 630 |
| *Oncorhynchus kisutch* | coho salmon | Salmoniformes | hypoxia inducible factor 1 subunit alpha, like 2 | HIF4A | hif1al2, npas1 | 109873637 | LG29 | 13 | XM_020465278 | XP_020320867 | 724 |
| *Oncorhynchus mykiss* | rainbow trout | Salmoniformes | hypoxia inducible factor 1 subunit alpha | HIF1Aa_s1 | hif1a | 100135944 | 19 | 15 | NM_001124288 | NP_001117760 | 766 |
| *Oncorhynchus mykiss* | rainbow trout | Salmoniformes | hypoxia-inducible factor 1-alpha-like | HIF1Aa_s2 | hif1al | 110505629 | 25 | 15 | XM_021584966 | XP_021440641 | 790 |
| *Oncorhynchus mykiss* | rainbow trout | Salmoniformes | endothelial PAS domain protein 1 | HIF2Aa_s1 | epas1 | 110499310 | 20 | 18 | XM_021576379 | XP_021432054 | 852 |
| *Oncorhynchus mykiss* | rainbow trout | Salmoniformes | endothelial PAS domain-containing protein 1-like | HIF2Aa_s2 | epas1b | 110503001 | 23 | 13 | XM_021581363 | XP_021437038 | 849 |
| *Oncorhynchus mykiss* | rainbow trout | Salmoniformes | endothelial PAS domain-containing protein 1-like | HIF2Ab | | 110492879 | 16 | 5 | XM_021567350 | XP_021423025 | 232 |
| *Oncorhynchus mykiss* | rainbow trout | Salmoniformes | hypoxia-inducible factor 3-alpha-like | HIF3A_s1 | hif3al | 110507593 | 27 | 15 | XM_021587678 | XP_021443353 | 676 |
| *Oncorhynchus mykiss* | rainbow trout | Salmoniformes | hypoxia-inducible factor 1-alpha-like | HIF3A_s2 | hif1al | 110503712 | 24 | 14 | XM_021582186 | XP_021437861 | 630 |
| *Oncorhynchus mykiss* | rainbow trout | Salmoniformes | hypoxia inducible factor 1 subunit alpha, like 2 | HIF4A | hif1al2 | 110510148 | 29 | 12 | XM_021591498 | XP_021447173 | 724 |
| *Oncorhynchus tshawytscha* | Chinook salmon | Salmoniformes | hypoxia inducible factor 1 subunit alpha | HIF1Aa_s1 | hif1a | 112261708 | 11 | 14 | XM_024437278 | XP_024293046 | 756 |
| *Oncorhynchus tshawytscha* | Chinook salmon | Salmoniformes | hypoxia-inducible factor 1-alpha-like | HIF1Aa_s2 | hif1al | 112256768 | 8 | 16 | XM_024430246 | XP_024286014 | 768 |
| *Oncorhynchus tshawytscha* | Chinook salmon | Salmoniformes | endothelial PAS domain protein 1 | HIF2Aa_s1 | epas1 | 112224583 | 25 | 16 | XM_024388216 | XP_024243984 | 853 |
| *Oncorhynchus tshawytscha* | Chinook salmon | Salmoniformes | endothelial PAS domain-containing protein 1-like | HIF2Aa_s2 | epas1b | 112260055 | 1 | 16 | XM_024434856 | XP_024290624 | 847 |
| *Oncorhynchus tshawytscha* | Chinook salmon | Salmoniformes | hypoxia-inducible factor 1 subunit alpha, like | HIF3A_s1 | hif1al | 112265295 | 13 | 14 | XM_024442447 | XP_024298215 | 572 |
| *Oncorhynchus tshawytscha* | Chinook salmon | Salmoniformes | hypoxia-inducible factor 1 subunit alpha, like 2 | HIF4A | hif1al2 | 112231158 | 33 | 12 | XM_024397745 | XP_024253513 | 724 |
| *Oreochromis niloticus* | Nile tilapia | Cichliformes | hypoxia inducible factor 1 subunit alpha | HIF1Aa | hif1a, hif1aa | 100703737 | LG19 | 15 | XM_005477038 | XP_005477095 | 769 |
| *Oreochromis niloticus* | Nile tilapia | Cichliformes | endothelial PAS domain protein 1b | HIF2Aa | epas1, epas1b | 100691768 | LG13 | 18 | XM_025897620 | XP_025753405 | 881 |
| *Oreochromis niloticus* | Nile tilapia | Cichliformes | endothelial PAS domain-containing protein 1 | HIF2Ab | epas1 | 100695204 | LG8 | 9 | XM_003441881 | XP_003441929 | 376 |
| *Oreochromis niloticus* | Nile tilapia | Cichliformes | hypoxia-inducible factor 1 subunit alpha, like | HIF3A | hif1al | 100708832 | LG14 | 15 | XM_005461141 | XP_005461198 | 679 |
| *Oryzias latipes* | Japanese ricefish | Beloniformes | hypoxia inducible factor 1 subunit alpha a | HIF1Aa | hif1aa | 101158737 | 22 | 15 | XM_023951295 | XP_023807063 | 750 |
| *Oryzias latipes* | Japanese ricefish | Beloniformes | endothelial PAS domain protein 1b | HIF2Aa | epas, epas1b | 101172602 | 15 | 16 | XM_023963599 | XP_023819367 | 861 |
| *Oryzias latipes* | Japanese ricefish | Beloniformes | endothelial PAS domain protein 1a | HIF2Ab | epas1a | 101168496 | 19 | 9 | XM_004080457 | XP_004080505 | 374 |

| Species | Common Name | Order | Genbank or Ensemble Gene Name | Group | Aliases | Gene ID (Genbank) | Chromosome # | Exons | mRNA Accession No. | Protein Accession No. | Protein Length (aa) |
|---|---|---|---|---|---|---|---|---|---|---|---|
| *Oryzias latipes* | Japanese ricefish | Beloniformes | hypoxia-inducible factor 1 subunit alpha, like | HIF3A | hif1al | 1011165163 | 13 | 16 | XM_004075255 | XP_004075303 | 650 |
| *Poecilia reticulata* | guppy | Cyprinodontiformes | hypoxia inducible factor 1 subunit alpha a | HIF1Aa | hif1aa | 103458897 | LG22 | 15 | XM_008400026 | XP_008398248 | 764 |
| *Poecilia reticulata* | guppy | Cyprinodontiformes | endothelial PAS domain-containing protein 1-like | HIF2Aa | epas1 | 103476630 | LG15 | 9 | XM_017308890 | XP_017164379 | 557 |
| *Poecilia reticulata* | guppy | Cyprinodontiformes | endothelial PAS domain protein 1a | HIF2Ab | epas1, epas1a | 103481538 | LG19 | 9 | XM_008437064 | XP_008435286 | 370 |
| *Poecilia reticulata* | guppy | Cyprinodontiformes | hypoxia-inducible factor 1 subunit alpha, like | HIF3A | hif1al | 103475214 | LG13 | 14 | XM_008426661 | XP_008424883 | 674 |
| *Salmo salar* | Atlantic salmon | Salmoniformes | hypoxia-inducible factor 1-alpha | HIF1Aa_s1 | hif1a | 106598919 | ssa01 | 15 | XM_014189950 | XP_014045425 | 762 |
| *Salmo salar* | Atlantic salmon | Salmoniformes | hypoxia-inducible factor 1-alpha-like | HIF1Aa_s2 | hif1al | 106610969 | ssa09 | 15 | XM_014210723 | XP_014066198 | 802 |
| *Salmo salar* | Atlantic salmon | Salmoniformes | endothelial PAS domain protein 1 | HIF2Aa_s1 | epas1 | 106589722 | ssa28 | 16 | XM_014179994 | XP_014035469 | 853 |
| *Salmo salar* | Atlantic salmon | Salmoniformes | endothelial PAS domain-containing protein 1-like | HIF2Aa_s2 | epas1 | 106611028 | ssa01 | 17 | XM_014210842 | XP_014066317 | 854 |
| *Salmo salar* | Atlantic salmon | Salmoniformes | hypoxia-inducible factor 1-alpha | HIF3A_s1 | hif1a | 100194993 | ssa20 | 16 | NM_001140022 | NP_001133494 | 628 |
| *Salmo salar* | Atlantic salmon | Salmoniformes | hypoxia-inducible factor 1-alpha-like | HIF3A_s2 | hif1al | 106612598 | ssa09 | 15 | XM_014213898 | XP_014069373 | 633 |
| *Salmo salar* | Atlantic salmon | Salmoniformes | hypoxia-inducible factor 1 subunit alpha, like 2 | HIF4A | hif1al2 | 106563688 | ssa11 | 12 | XM_014129489 | XP_013984964 | 725 |
| *Salvelinus alpinus* | arctic char | Salmoniformes | hypoxia-inducible factor 1-alpha | HIF1Aa_s1 | hif1a | 111968658 | LG9 | 14 | XM_023994425 | XP_023850193 | 756 |
| *Salvelinus alpinus* | arctic char | Salmoniformes | hypoxia-inducible factor 1-alpha | HIF1Aa_s2 | hif1a | 111963342 | LG4q.2 | 16 | XM_023986707 | XP_023842475 | 759 |
| *Salvelinus alpinus* | arctic char | Salmoniformes | endothelial PAS domain protein 1 | HIF2Aa_s1 | epas1 | 111968049 | LG8 | 17 | XM_023993573 | XP_023849341 | 853 |
| *Salvelinus alpinus* | arctic char | Salmoniformes | endothelial PAS domain-containing protein 1-like | HIF2Aa_s2 | epas1l | 112073362 | Un | 15 | XM_024140674 | XP_023996442 | 849 |
| *Salvelinus alpinus* | arctic char | Salmoniformes | hypoxia-inducible factor 1-alpha | HIF3A_s1 | hif1a | 111982824 | LG22 | 15 | XM_024014433 | XP_023870201 | 629 |
| *Salvelinus alpinus* | arctic char | Salmoniformes | hypoxia-inducible factor 1-alpha | HIF3A_s2 | hif1a | 111949752 | LG22 | 16 | XM_023967103 | XP_023822871 | 659 |
| *Salvelinus alpinus* | arctic char | Salmoniformes | hypoxia inducible factor 1 subunit alpha, like 2 | HIF4A | hif1al2, npas1 | 111964361 | LG5 | 12 | XM_023988241 | XP_023844009 | 721 |
| *Scleropages formosus* | Asian arowana | Osteoglossiformes | hypoxia inducible factor 1 subunit alpha a | HIF1Aa | hif1a, hif1aa | 108919562 | 15 | 15 | XM_018727619 | XP_018583135 | 762 |
| *Scleropages formosus* | Asian arowana | Osteoglossiformes | endothelial PAS domain protein 1b | HIF2Aa | epas1, epas1b | 108942753 | 8 | 18 | XM_018766230 | XP_018621746 | 881 |
| *Scleropages formosus* | Asian arowana | Osteoglossiformes | hypoxia inducible factor 1 subunit alpha, like | HIF3A | hif1al | 108928738 | 10 | 19 | XM_018742811 | XP_018598327 | 628 |
| *Scleropages formosus* | Asian arowana | Osteoglossiformes | hypoxia inducible factor 1 subunit alpha, like 2 | HIF4A | hif1al2 | 108934068 | 4 | 15 | XM_018751566 | XP_018607082 | 711 |
| *Takifugu rubripes* | torafugu | Tetraodontiformes | hypoxia inducible factor 1 subunit alpha a | HIF1Aa | hif1a, hif1aa | 101071027 | 2 | 15 | XM_003962474 | XP_003962523 | 755 |
| *Takifugu rubripes* | torafugu | Tetraodontiformes | endothelial PAS domain protein 1b | HIF2Aa | epas1, epas1b | 101067536 | 4 | 16 | XM_011603052 | XP_011601354 | 854 |
| *Takifugu rubripes* | torafugu | Tetraodontiformes | endothelial PAS domain protein 1 | HIF2Ab | epas1 | 101073099 | 1 | 9 | XM_003976888 | XP_003976937 | 359 |
| *Takifugu rubripes* | torafugu | Tetraodontiformes | hypoxia inducible factor 1 subunit alpha, like | HIF3A | hif1al | 101062698 | 11 | 16 | XM_029843274 | XP_029699134 | 676 |
| *Xiphophorus maculatus* | southern platyfish | Cyprinodontiformes | hypoxia inducible factor 1 subunit alpha | HIF1Aa | hif1aa | 102217674 | 19 | 15 | XM_023352592 | XP_023208360 | 758 |
| *Xiphophorus maculatus* | southern platyfish | Cyprinodontiformes | endothelial PAS domain protein 1b | HIF2Aa | epas1, epas1b | 102218651 | 22 | 16 | XM_023326963 | XP_023182731 | 867 |
| *Xiphophorus maculatus* | southern platyfish | Cyprinodontiformes | endothelial PAS domain protein 1a | HIF2Ab | epas1, epas1a | 102227739 | 10 | 9 | XM_005794777 | XP_005794834 | 409 |
| *Xiphophorus maculatus* | southern platyfish | Cyprinodontiformes | hypoxia inducible factor 1 subunit alpha, like | HIF3A | hif3al, hif1al2 | 102223582 | 18 | 15 | XM_005811977 | XP_005812034 | 650 |

| Order | Species | Gene | -10 | -9 | -8 | -7 | -6 | -5 | -4 | -3 | -2 | -1 | 0 | 1 | 2 | 3 | 4 | 5 | 6 | 7 | 8 | 9 | 10 | Gene ID | Chr/LG | mRNA Accession | Protein Accession |
|---|---|---|---|---|---|---|---|---|---|---|---|---|---|---|---|---|---|---|---|---|---|---|---|---|---|---|---|
| Lepisosteiformes | *Lepisosteus oculatus* | HIF1A | pcnxl4-> | dhrs7<- | ppm1a-> | six6-> | six1<- | six4<- | mnat1-> | trmt5<- | slc38a6-> | prkch-> | **hif1a->** | snapc1-> | syt16-> | kcnh5<- | rhoj-> | gphb5<- | ppp2r5e<- | wdr89<- | sgpp1<- | syne2-> | esr2a<- | 102682513 + 107077742 | LG 7 | XM_015350184 + XM_015350128 | XP_015205670 + XP_015205614 |
| Osteoglossiformes | *Scleropages formosus* | HIF1Aa | pcnxl4-> | dhrs7<- | ppm1a-> | six6-> | six1<- | six4<- | mnat1-> | trmt5<- | slc38a6-> | prkch-> | **hif1aa->** | snapc1-> | syt16-> | sgpp1-> | wdr89-> | ppp2e5e-> | inf2-> | kcnh5<- | gphb5<- | syne2-> | esr2<- | 108919562 | Chr 15 | XM_018727619 | XP_018583135 |
| Cypriniformes | *Danio rerio* | HIF1Aa | pcnxl4-> | dhrs7<- | ppm1aa-> | six6a<- | six1a<- | six4a<- | mnat1-> | trmt5<- | slc38a6-> | prkcha-> | **hif1aa->** | unknown<- | sertad4<- | syt14a<- | diexf<- | osr1-> | nt5c1ba-> | rdh14a-> | kcns3a<- | wdr35<- | matn3b<- | 797150 | Chr 13 | NM_001308559 | NP_001295488 |
| Characiformes | *Astyanax mexicanus* | HIF1Aa | pcnx4-> | dhrs7<- | ppm1a-> | six6<- | six1b<- | tchhl<- | mnat1-> | trmt5<- | slc38a6-> | prkch-> | **hif1a->** | unknown<- | sertad4<- | syt14<- | diexf<- | osr1-> | ccdc106<- | rdh14-> | kcns3<- | wdr35<- | matn3b<- | 103022448 | Chr 25 | XM_007247542 | XP_007247604 |
| Esosiformes | *Esox lucius* | HIF1Aa | pcnx4-> | dhrs7<- | ppm1a-> | six6-> | six1<- | six4<- | mnat1-> | trmt5<- | slc38a6-> | prkcha-> | **hif1aa->** | snapc1-> | syt16-> | sgpp1-> | wdr89-> | ppp2e5e-> | kcnh5-> | gphb5<- | syne2-> | esr2b<- | jagged-2<- | 105015558 | Chr 15 | NM_001310847 | NP_001297776 |
| Salmoniformes | *Oncorhynchus mykiss* | HIF1Aa_s1 | pcnx4-> | unknown-> | ppm1a-> | six6-> | six1<- | six4<- | mnat1-> | trmt5<- | slc38a6-> | prkcha-> | **hif1a->** | snapc1-> | syt16-> | sgpp1<- | ppp2r5e<- | kcnh5-> | gphb5<- | nesprin-2-like-> | titan homolog-> | syne2-> | esr2a<- | 100135944 | Chr 19 | NM_001124288 | NP_001117760 |
| Salmoniformes | *Oncorhynchus mykiss* | HIF1Aa_s2 | rtn1<- | lrrc9-> | unknown-> | dhrs7<- | ppm1a-> | six1<- | six4<- | mnat-> | slc38a6-> | prkch-> | **hif1al->** | sacsin-like<- | znf239-> | wdr89-> | ppp2r5eb-> | gphb5<- | esr2<- | jagged-1b-like<- | unknown<- | unknown-> | brf1a<- | 110505629 | Chr 25 | XM_021584966 | XP_021440641 |
| Cyprinodontiformes | *Xiphophorous maculatus* | HIF1Aa | pcnx4-> | dhrs7<- | ppm1a-> | six6-> | six1<- | six4<- | mnat1-> | trmt5<- | slc38a6-> | prkch-> | **hif1aa->** | htra2<- | mis18bp1-> | tpp1-> | yy1<- | degs2-> | evl<- | eml1<- | unknown<- | unknown-> | pkdcc<- | 102217674 | Chr 19 | XM_023352592 | XP_023208360 |
| Tetraodontiformes | *Takifugu rubripes* | HIF1Aa | dhrs7<- | ppm1a-> | six6-> | ino80c-> | six1<- | six4<- | mnat1-> | trmt5<- | slc38a6-> | prkch-> | **hif1a->** | htra2<- | mis18bp1-> | tpp1-> | yy1a<- | degs2-> | evl<- | eml1<- | pkdcc<- | disp2-> | rpusd2<- | 101071027 | Chr 2 | XM_003962474 | XP_003962523 |
| Cypriniformes | *Danio rerio* | HIF1Ab | ppp1r13bb<- | atp5mpl<- | rd3l<- | rtn1b<- | ppm1ab-> | six6b-> | six1b<- | six4b<- | unknown<- | prkchb-> | **hif1ab->** | snapc1a-> | sgpp1-> | ppp2r5ea-> | rhoj<- | kcnh5a-> | ptk2bb-> | chrna2b<- | unknown<- | baalc-> | nme6<- | 393292 | Chr 20 | NM_001310042 | NP_001296971 |
| Characiformes | *Astyanax mexicanus* | HIF1Ab | ckb-> | ppp1r13b<- | atp5mpl<- | rd3l<- | rtn1b<- | ppm1ab-> | six6-> | six1b<- | six4<- | prkch-> | **hif1ab->** | snapc1a-> | sgpp1-> | ppp2r5ea-> | rhoj<- | kcnh5-> | ptk2b-> | chrna2<- | cstl-> | baal-> | nme6<- | 103033873 | Chr 3 | XM_007256662 | XP_007256724 |
| **Order** | **Species** | **Gene** | **-10** | **-9** | **-8** | **-7** | **-6** | **-5** | **-4** | **-3** | **-2** | **-1** | **0** | **1** | **2** | **3** | **4** | **5** | **6** | **7** | **8** | **9** | **10** | **Gene ID** | **Chr/LG** | **mRNA Accession** | **Protein Accession** |
| Lepisosteiformes | *Lepisosteus oculatus* | HIF2A | abcg5<- | lrpprc<- | ppm1b-> | slc3a1-> | prepl<- | camkmt-> | six3-> | six2<- | srdb1<- | prkce-> | **epas1b->** | rhoq-> | pigf<- | cript-> | gch2<- | socs5-> | prop1-> | mcfd2<- | ttc7a-> | clg16h2orf61<- | calm2<- | 102694568 | LG 16 | XM_015362793 | XP_015218279 |
| Osteoglossiformes | *Scleropages formosus* | HIF2Aa | abcg8<- | lrpprc<- | ppm1b-> | slc3a1-> | prepl<- | camkmt-> | six3-> | six2<- | srdb1<- | prkce-> | **epas1b->** | rhoq-> | gch1<- | socs5-> | prop1-> | mcfd2<- | ttc7a-> | calm2<- | epcam-> | kcnk12<- | msh6-> | 108942753 | Chr 8 | XM_018766230 | XP_018621746 |
| Cypriniformes | *Danio rerio* | HIF2Aa | plekhh2-> | epcam<- | fbn2-> | calm2a-> | clg16h2orf61-> | ttc7a<- | mcfd2-> | prop1<- | socs5b<- | prkceb-> | **epas1b->** | dhx57-> | galm<- | atl2-> | unknown-> | hells<- | unknown-> | prkg2<- | col13Aa<- | h2afy2-> | gbf1<- | 555192 | Chr 13 | NM_001039806 | NP_001034895 |
| Characiformes | *Astyanax mexicanus* | HIF2Aa | prop1-> | mcfd2<- | ttc7a-> | stpg4<- | calm3<- | fbn1<- | epcam-> | plekhh2<- | thada-> | prkce-> | **epas1b->** | srdb1-> | six2-> | six3<- | camkmt<- | prepl-> | slc3a1<- | ppm1b<- | lrpprc-> | abcg8<- | abcg5-> | 103026643 | Chr 25 | XM_022684458 | XP_022540179 |
| Esosiformes | *Esox lucius* | HIF2Aa | abcg8-> | lrpprc<- | ppm1ba-> | slc3a1-> | prepl<- | camkmt-> | six3a-> | six2a<- | srbd1<- | prkce-> | **epas1b->** | unknown<- | pcgf5a<- | ankrd1-> | mertka-> | tmem87b-> | fbln7<- | zc3h6-> | unknown<- | itga9<- | golga4<- | 105021253 | Chr 5 | XM_010888790 | XP_010887092 |
| Salmoniformes | *Oncorhynchus mykiss* | HIF2Aa_s1 | abcg8-> | lrpprc<- | ppm1b-> | slc3a1-> | prepl<- | camkmt-> | six3-> | six2<- | srdb1<- | prkce-> | **epas1->** | pcbf5b<- | ankrd1a-> | lhr-> | mertka-> | tmem87b-> | fbln7<- | zc3h6-> | unknown<- | itga9<- | golga4<- | 110499310 | Chr 20 | XM_021576379 | XP_021432054 |
| Salmoniformes | *Oncorhynchus mykiss* | HIF2Aa_s2 | perp<- | arfgef3-> | catspere<- | adss2-> | zbtb18<- | cep170aa-> | dync2li1-> | six3a-> | six2a<- | prkce-> | **epas1l->** | prkce<- | pcgf5a<- | mertka-> | tmem87al-> | fbln7<- | itga9<- | golga4<- | kcnk12-> | fbxo11a<- | foxn2<- | 110503001 | Chr 23 | XM_021581363 | XP_021437038 |
| Cyprinodontiformes | *Xiphphorous maculatus* | HIF2Aa | itsn1<- | setd4<- | cbr1-> | nrg3<- | oard1-> | unknown<- | ccdc172<- | gfra1-> | atrnl1<- | prkce-> | **epas1->** | unknown<- | cox20<- | cgbb-> | ndst2<- | zswim8<- | polr1c -> | fbxo28 -> | degs1-> | nvl<- | spata2l-> | 102218651 | Chr 22 | XM_023326963 | XP_023182731 |
| Tetraodontiformes | *Takifugu rubripes* | HIF2Aa | itsn1<- | cryzl1-> | setd4<- | cbr1-> | nrg3-> | unknown-> | ccdc172<- | gfra1a-> | atrnl1a<- | prkce-> | **epas1->** | slc22a7-> | slc5a7l-> | cox20<- | cgbb-> | nckap1<- | dusp19-> | unknown-> | zc3h15-> | rbm45<- | cyc-> | 101067536 | Chr 4 | XM_011603052 | XP_011601354 |
| Cypriniformes | *Danio rerio* | HIF2Ab | rhoq<- | cox7a3<- | kcng3<- | mta3-> | hcar1-4<- | zfp36l2<- | ppm1bb-> | six3b-> | six2b<- | prkcea-> | **epas1a->** | glud1b-> | sncgb<- | mmrn2b-> | bmpr1ab<- | ldb3b<- | opn4b<- | usp54b<- | myoz1b<- | synpo2lb<- | itgb3b-> | 566886 | Chr 12 | XM_690170 | XP_695262 |
| Characiformes | *Astyanax mexicanus* | HIF2Ab | cox7a3<- | kcng3<- | mta3-> | hcar1-4<- | unknown<- | zfp36l2<- | ppm1b-> | six3-> | six2<- | prkce-> | **epas1->** | glud1-> | sncgb<- | mmrn2l-> | bmpr1a<- | ldb3<- | opn4a<- | unknown-> | usp54<- | myoz1<- | synpo2l<- | 103036701 | Chr 6 | XM_022670333 | XP_022526054 |
| Salmoniformes | *Oncorhynchus mykiss* | HIF2Ab | tbcc-> | prph2-> | qki-a-> | timm23-> | unknown<- | unknown<- | unknown<- | trim21<- | prss12-> | camk2d<- | **epas1l->** | glud1-> | snca<- | opn4a<- | unknown<- | usp54<- | myoz1<- | synpo2l<- | sec24c-> | nlrp12<- | nrnx1a<- | 110492879 | Chr 16 | XM_021567350 | XP_021423025 |
| Cyprinodontiformes | *Xiphphorous maculatus* | HIF2Ab | unknown<- | gch2-> | cript<- | pigf-> | cox7al<- | kcng3<- | mta3-> | ppm1b-> | six3-> | prkce-> | **epas1->** | glud1-> | sncgb<- | ldb3b<- | opn4a<- | usp54<- | myoz1<- | synpo2l<- | sec24c-> | nlrp3<- | camk2g-> | 102227739 | Chr 10 | XM_005794777 | XP_005794834 |
| Tetraodontiformes | *Takifugu rubripes* | HIF2Ab | msh2<- | gch1-> | cript<- | pigf-> | cox7al<- | kcng3<- | mta3-> | ppm1b-> | six3-> | prkce-> | **epas1->** | glud1-> | | | | | poor or missing annotation | | | | | 101073099 | Chr 1 | XM_003976888 | XP_003976937 |
| **Order** | **Species** | **Gene** | **-10** | **-9** | **-8** | **-7** | **-6** | **-5** | **-4** | **-3** | **-2** | **-1** | **0** | **1** | **2** | **3** | **4** | **5** | **6** | **7** | **8** | **9** | **10** | **Gene ID** | **Chr/LG** | **mRNA Accession** | **Protein Accession** |
| Lepisosteiformes | *Lepisosteus oculatus* | HIF3A | prss16<- | hnrnpl<- | srsf7a<- | slc8a2<- | unknown<- | actn4<- | eif3k<- | map4k1-> | wdr20-> | tm7sf2l-> | **hif1al->** | nccrp1<- | susd6l<- | pak4<- | ankrd34a-> | ccdc9<- | unknown-> | sae1<- | zc3h4-> | cdc42ep2l-> | ercc2-> | 102689100 | LG 2 | XM_015340661 | XP_015196147 |
| Osteoglossiformes | *Scleropages formosus* | HIF3A | hnrnpl<- | srsf7a<- | slc8a2<- | smoc1<- | actn4<- | capn9<- | eif3k<- | map4k-> | wdr20-> | tm7sf2l-> | **hif1al->** | susd6l<- | pak4<- | ccdc9<- | bbc3-> | sae1<- | zc3h4-> | cdc42ep2l-> | ercc2-> | klc3<- | ckm-> | 108928738 | Chr 10 | XM_018742811 | XP_018598327 |
| Cypriniformes | *Danio rerio* | HIF3A | rpa1-> | rtn4rl1b<- | dph1-> | hic1<- | tsr1<- | taok1a-> | tp53inp13-> | mmp20b-> | mmp20a-> | npat<- | **hif1al->** | susd6l<- | pak4<- | aifm4<- | tlcd2-> | ccdc92-> | cluha-> | pafah1b1a-> | mettl16-> | mnta-> | slc35f2l<- | 393376 | Chr 15 | XM_005157571 | XP_005157628 |
| Characiformes | *Astyanax mexicanus* | HIF3A | rpa1-> | rtn4rl1b<- | dph1-> | hic1<- | tsr1<- | taok1-> | tp53i13-> | mmp20-> | mmp20-> | npat<- | **hif1al->** | susd6<- | pak4<- | aifm3<- | tlcd2-> | ccdc92b-> | cluh-> | pafah1b1a-> | mettl16-> | mnt-> | slc35f2<- | 103027586 | Chr 1 | XM_007245906 | XP_007245968 |
| Esosiformes | *Esox lucius* | HIF3A | hnrnpl<- | srsf7<- | slc8a2b<- | smoc1<- | actn4<- | capn3<- | unknown<- | eif3k<- | map4k5-> | wdr20-> | **hif1al->** | susd6<- | pak4<- | ccdc9<- | bbc3-> | sae1<- | zc3h4-> | cdc42ep2-> | ercc2-> | klc3<_ | mark4a<- | 105012786 | Chr 1 | XM_029120020 | XP_028975853 |
| Salmoniformes | *Oncorhynchus mykiss* | HIF3A_s1 | prss16<- | ech1<- | hnrnpl<- | srsf7<- | slc8a2<- | smoc1<- | actn4<- | eif3k<- | map4k5l-> | wdr20-> | **hif3al->** | susd6l<- | pak4<- | ccdc9<- | sae1<- | zc3h4l-> | cdc42ep2-> | ercc2-> | klc3<- | ckm-> | mark4<- | 110507593 | Chr 27 | XM_021587678 | XP_021443353 |
| Salmoniformes | *Oncorhynchus mykiss* | HIF3A_s2 | meis3<- | prss16<- | hnrnpl<- | srsf7<- | slc8a2b<- | actn4<- | capn9l<- | eif3k<- | map4k5l-> | wdr20-> | **hif1al->** | susd6l<- | pak4<- | ccdc9<- | sae1<- | zc3h4l-> | cdc42ep2-> | klc3<- | ckm-> | mark4<- | igfals-> | 110503712 | Chr 24 | XM_021582186 | XP_021437861 |
| Cyprinodontiformes | *Xiphphorous maculatus* | HIF3A | vstm5<- | unknown-> | hnrnpl<- | srsf7<- | slc8a2<- | eik3k<- | capn9<- | actn4<- | map4k5l-> | wdr20-> | **hif3al->** | susd6l<- | pak4<- | ccdc9<- | sae1<- | zc3h4l-> | cdc42ep2-> | ercc2-> | srcrm130<- | klc3<- | ckm-> | 102223582 | Chr 18 | XM_005811977 | XP_005812034 |
| Tetraodontiformes | *Takifugu rubripes* | HIF3A | gemin4-> | fam57a<- | hnrnpl<- | srsf7<- | slc8a2<- | actn4<- | capn9l<- | eif3k<- | map4k5l-> | wdr20-> | **hif1al->** | susd6l<- | pak4<- | ccdc9<- | unknown-> | sae1<- | zc3h4l-> | cdc42ep2-> | ercc2-> | klc3<- | ckm-> | 101062698 | Chr 11 | XM_029843274 | XP_029699134 |
| **Order** | **Species** | **Gene** | **-10** | **-9** | **-8** | **-7** | **-6** | **-5** | **-4** | **-3** | **-2** | **-1** | **0** | **1** | **2** | **3** | **4** | **5** | **6** | **7** | **8** | **9** | **10** | **Gene ID** | **Chr/LG** | **mRNA Accession** | **Protein Accession** |
| Lepisosteiformes | *Lepisosteus oculatus* | HIF4A | slc22a6<- | capn1l<- | klc2<- | pacs1<- | zdhhc24-> | catsper1<- | prkd3l-> | actn3<- | ccs<- | dpp3<- | **epas1l->** | six6l<- | b3gnt1<1 | pfdn2<- | nit1-> | col3a1l-> | sstr1l-> | cd248<- | rce1<- | peli3l-> | rab1bl<- | 102687670 | LG 28 | XM_015338767 | XP_015194253 |
| Osteoglossiformes | *Scleropages formosus* | HIF4A | cfb-> | bbs1<- | capn1<- | sf3b2-> | pacs1-> | actn3<- | gal3st3-> | rin1a-> | znf664-> | dpp3<- | **hif1al2->** | six6l<- | pfdn2<- | sstr1l-> | cd248<- | rce1<- | peli3-> | rab1bl<- | muc17<- | fam89b<- | zrdn2<- | 108934068 | Chr 4 | XM_018751566 | XP_018607082 |
| Cypriniformes | *Danio rerio* | HIF4A | capn1<- | bbs1<- | cfbl<- | bfb-> | cfb<- | slc29a2-> | actn3a<- | rbm14a<- | rin1a-> | dpp3<- | **hif1al2->** | sart1<- | b3gnt2l<- | dusp10<- | mark2a-> | unknown-> | unknown-> | bada<- | slc8a4a-> | ppp1r14ba-> | fkbp2<- | 497283 | Chr 21 | NM_001012371 | NP_001012371 |
| Characiformes | *Astyanax mexicanus* | HIF4A | mrps17<- | fibh-> | limk1-> | unknown<- | unknown-> | cfb-> | slc29a2-> | rbm4b<- | rin2-> | dpp3<- | **hif1al2->** | sart1<- | b3gnt2<- | unknown<- | mark2-> | lark-> | bada<- | slc8a1-> | ppp1r14b-> | fkbp2<- | pc<- | 103041845 | Chr 14 | XM_007244478 | XP_007244540 |
| Esosiformes | *Esox lucius* | HIF4A | cfbl-> | bbs1-> | capn1<- | cfb<- | pacs1a-> | actn3a<- | rbm14a<- | rin1a-> | dpp3<- | il2rg<- | **hif1al2->** | cd248b<- | rab1bb<- | ehbp1l1b<- | znrd2<- | slc35f4<- | arr3a<- | mcf2-> | f9b<- | fgf13b-> | cldn4-> | 105024933 | Chr 7 | XM_010895238 | XP_010893540 |
| Salmoniformes | *Oncorhynchus mykiss* | HIF4A | ccnb3l<- | tnfsf10l-> | clcn5l<- | leap2b-> | picalm1<- | rab39b-> | slc7a3a<- | snx12l<- | il2rg<- | unknown-> | **hif1al2->** | cd248b<- | rab1al<- | ehbp1l1<- | fam89b<- | sssca1<- | slc35f3<- | arrc<- | fgf13b-> | zic3-> | tmpops3b<- | 110510148 | Ch 29 | XM_021591498 | XP_021447173 |

| Gene Group | Fish Group | Count | Exons (MIN) | Exons (MAX) | Exons (MEDIAN) | Amino Acids (MIN) | Amino Acids (MAX) | Amino Acids (MEDIAN) |
|---|---|---|---|---|---|---|---|---|
| HIF1Aa | Asian arowana | 1 | 15 | 15 | 15 | 762 | 762 | 762 |
| HIF1Aa | Otocephala | 5 | 14 | 15 | 15 | 611 | 748 | 717 |
| HIF1Aa_s1 | Salmoniformes | 5 | 14 | 15 | 14 | 756 | 766 | 762 |
| HIF1Aa_s2 | Salmoniformes | 5 | 15 | 16 | 15 | 758 | 802 | 768 |
| HIF1Aa | Neoteleostei | 9 | 15 | 16 | 15 | 748 | 774 | 758 |
| HIF1Ab | Otocephala | 5 | 14 | 16 | 15 | 774 | 798 | 777 |
| HIF2Aa | Asian arowana | 1 | 18 | 18 | 18 | 881 | 881 | 881 |
| HIF2Aa | Otocephala | 5 | 10 | 17 | 16 | 565 | 912 | 834 |
| HIF2Aa_s1 | Salmoniformes | 5 | 16 | 18 | 16 | 852 | 853 | 853 |
| HIF2Aa_s2 | Salmoniformes | 5 | 13 | 17 | 16 | 845 | 854 | 849 |
| HIF2Aa | Neoteleostei | 9 | 9 | 18 | 16 | 557 | 881 | 860 |
| HIF2Ab | Otocephala | 5 | 10 | 17 | 16 | 459 | 845 | 816 |
| HIF2Ab | Salmoniformes | 2 | 5 | 9 | 7 | 232 | 391 | 312 |
| HIF2Ab | Neoteleostei | 8 | 8 | 9 | 9 | 359 | 409 | 370 |
| HIF3A | Asian arowana | 1 | 19 | 19 | 19 | 628 | 628 | 628 |
| HIF3A | Otocephala | 5 | 15 | 17 | 15 | 626 | 646 | 633 |
| HIF3A_s1 | Salmoniformes | 5 | 14 | 16 | 15 | 572 | 676 | 628 |
| HIF3A_s2 | Salmoniformes | 4 | 14 | 16 | 15 | 630 | 659 | 632 |
| HIF3A | Neoteleostei | 9 | 12 | 16 | 15 | 649 | 679 | 662 |
| HIF4A | Asian arowana | 1 | 15 | 15 | 15 | 711 | 711 | 711 |
| HIF4A | Otocephala | 5 | 14 | 15 | 14 | 606 | 708 | 663 |
| HIF4A | Salmoniformes | 5 | 12 | 13 | 12 | 721 | 725 | 724 |
